# High-Throughput Centrifuge Force Microscopy Reveals Dynamic Immune-Cell Avidity at the Single-Cell Level

**DOI:** 10.1101/2025.02.27.640408

**Authors:** Hans T. Bergal, Koji Kinoshita, Wesley P. Wong

**Author notes:** Correspondence: Wesley P. Wong, Center for Life Sciences, 3rd floor, Boston Children’s Hospital, 3 Blackfan Circle, Boston, MA 02215. These authors contributed equally to this work (Hans T. Bergal, Koji Kinoshita).

## Abstract

Cell-cell binding, mediated by the physical interactions of receptors and their ligands, plays a fundamental role in immune processes such as immune surveillance and T-cell activation. However, current approaches for measuring cell avidity often lack either throughput or quantitative precision. Here, we introduce a high-throughput approach for quantifying cell binding lifetimes and strength using a centrifuge force microscope (CFM)—a compact microscope operating within a standard benchtop centrifuge. The CFM enables live monitoring of single-cell interactions under force, conducting thousands of force experiments in parallel. To facilitate the real-time study of live cell interactions, we developed a next-generation CFM with multichannel fluorescence imaging capabilities. This system accommodates measurements in two modes: cell-protein binding and cell-cell avidity assays. Using this system, we investigated immune-cell binding mediated by Bispecific T-cell Engager (BiTE) molecules, a novel immunotherapy designed to enhance immune-cell targeting of cancer cells. In cell-protein assays, we quantified T- and B-cell unbinding from BiTE-functionalized surfaces, revealing receptor-specific relationships between ligand concentration and binding strength. In cell-cell assays, we examined BiTE-mediated binding of T-cells to Nalm6 B-cells, a precursor leukemia cell line, uncovering a strong, time-dependent increase in BiTE-mediated immune-cell avidity. By integrating high-throughput and quantitative single-cell force analysis, the CFM provides new insights into the dynamic nature of immunological interactions under force, with broad implications for immunotherapy and cellular mechanics.

## 1. Introduction

Cell-cell binding is fundamental to many biological processes, from immune sensing to neuronal development. Beyond mere attachment, these interactions can exert mechanical forces, leading to changes in cytoskeletal networks, opening of ion channels, and activation of signaling pathways^1, 2, 3, 4^. While many methods have been developed to measure the thermodynamic and kinetic parameters of individual protein-protein interactions^5^, the interaction of single isolated proteins in solution may not directly translate to how they behave on the cell surface in coordination with other molecules, particularly under force^6, 7^.

A wide range of parameters, such as the identities of the proteins involved, the number of interactions, their spatial distribution, and time dynamics, can all contribute to the interaction strength between cells^7, 8^. For example, adhesion between an antigen-presenting cell (APC) and a T-cell evolves from a weak interaction between a T-cell receptor (TCR) and major histocompatibility complex (MHC) proteins into a strong interaction composed of integrins organized into an immune synapse^8^. T-cell responses can be further influenced by the affinity of TCRs^9^, the number of receptors^10^, and the applied force vector^4^. Measuring cellular avidity in reaction to drugs, cell contacts, applied force, or over time would allow us to probe cellular behavior and gain critical insights into mechanisms of cell interaction and adaptation.

Cell-cell interactions often exhibit strong binding interactions, with lifetimes extending beyond feasible experimental timeframes^11^. Moreover, the dynamic nature of the interactions that govern cell-cell adhesion requires precise, time-resolved measurements of off-rate kinetics to capture their behavior accurately^12^. The mechanical force of cell adhesion has also been shown to play a critical role in immune processes like T-cell activation, proliferation, and cytotoxicity^2^. Recent force spectroscopy has revealed the importance of force in T-cell receptor specificity^13^ and demonstrated long lifetimes under physiological loads^10^. These findings underscore the importance of applying mechanical force to better understand immune cell behavior.

Over the past decades, micromanipulation techniques—including atomic force microscopy (AFM)^14^, optical tweezers^13^, and the biomembrane force probe (BFP)^15, 16^—as well as multiplexed systems such as hydrodynamic flow setups^17^ and acoustic force spectroscopy (AFS)^18, 19^, have been successfully used to measure cell avidity^20, 21^. While micromanipulation methods offer high temporal and force resolution, they often measure cells one at a time, which slows data collection, limits the characterization of cell heterogeneity, and hinders their broader adoption^22, 23, 24^. AFS and hydrodynamic flow systems enable parallel measurements but generally produce non-uniform force fields, require external calibration due to complex input-to-force mapping, and may exert unwanted torque on cells.

To increase the throughput, accuracy, and accessibility of cell force spectroscopy, we employed the centrifuge force microscope (CFM), a specialized instrument designed to simultaneously apply centrifugal force to interrogate cell adhesion^25^. The CFM, a microscope integrated into a benchtop centrifuge, combines real-time imaging with straightforward force quantification during centrifugation^26^. Its use of high-resolution imaging and a uniform centrifugal force field enables consistent, parallel measurements across the entire field of view^27, 28^. In one of its earliest applications, molecules were tethered between a surface and beads, enabling the simultaneous characterization of hundreds to thousands of single-molecule interactions under force, thereby expanding the achievable throughput of single-molecule force spectroscopy^25, 26^.

Here, we introduce a next-generation multichannel fluorescence CFM to quantify both cell-protein and cell-cell interactions under force (**Figure 1A)**. Cell-protein interactions are quantified by attaching proteins of interest to a surface, allowing cells to bind, and then measuring bond lifetimes by monitoring the real-time response of each cell under applied centrifugal force. To extend this approach, we substitute the protein-functionalized surface with a cell monolayer, enabling the direct measurement of cell-cell binding strength. Measuring cell-cell avidity with the CFM allows us to examine complex interactions under physiological receptor densities without requiring labor-intensive protein purification. Applying force enables the probing of cell avidity strength throughout the progression of cell interactions, from initial weak binding events—such as TCR-MHC engagement—to the formation of a stable immune synapse. This approach reveals dynamic insights that traditional methods like sequencing or proximity-based chemical tags often miss^29^, and addresses the inherent difficulty of studying dynamic, force-dependent cell-cell interactions in their natural context.

**Figure 1.**
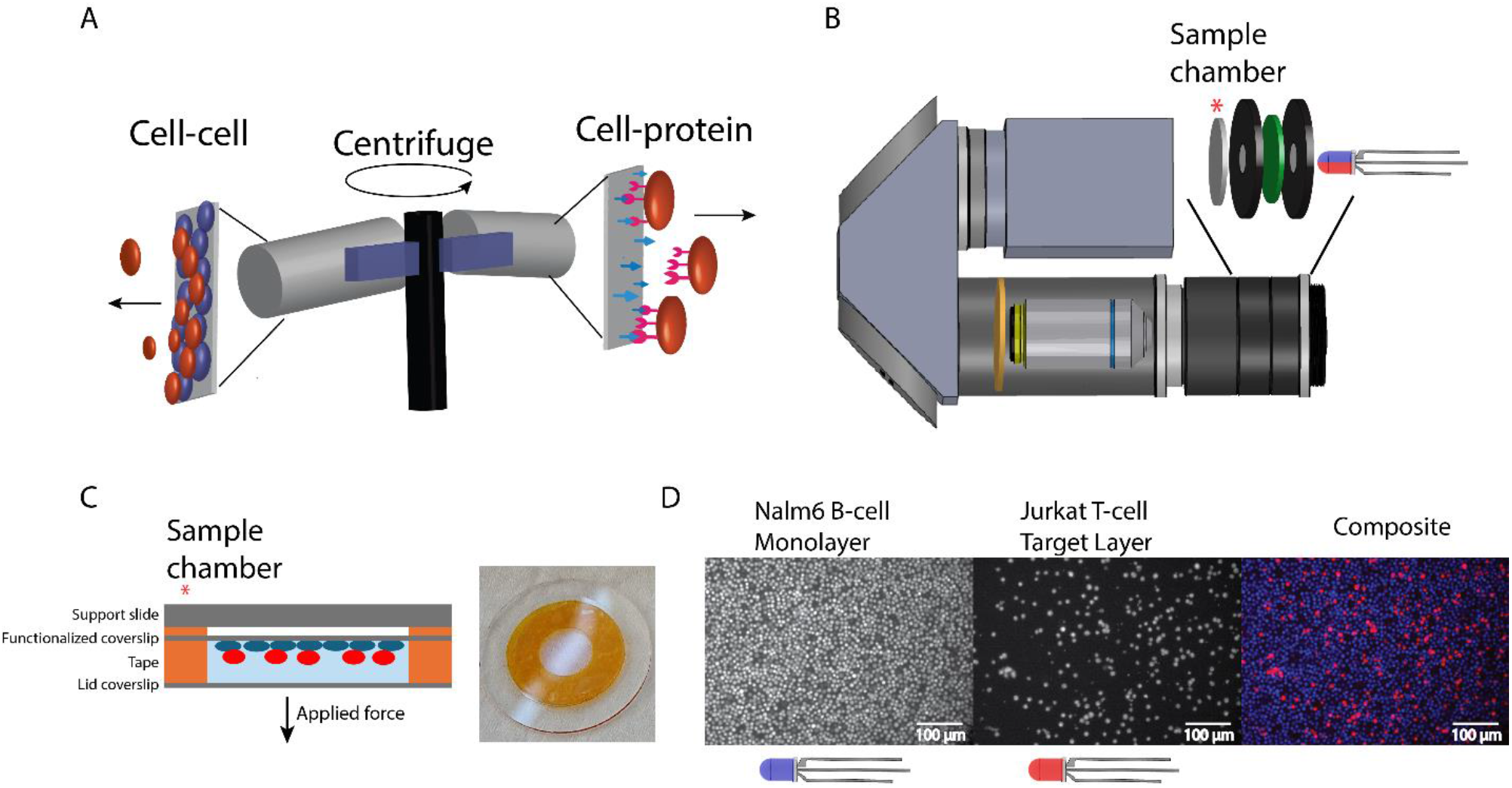
Overview of making cell-avidity measurements with the fluorescence centrifuge force microscope (CFM). **(A)** This schematic shows the measurement setup, where target cells (red) are added onto a monolayer (blue) or a surface functionalized with the target molecule. Centrifugal force is applied, causing unbound target cells to detach. **(B)** The fluorescence CFM setup updates previous designs with additional components that include an excitation filter (green), emission filter (yellow), and multicolor LED. **(C)** The sample chamber consists of two glass pieces held together with double-sided tape and an additional support slide. Cells attach to one side with poly-L-lysine (PLL) before being loaded into the fluorescence CFM. **(D)** Fluorescent images display labeled monolayer Nalm6 B-cells and Jurkat T-cells captured with the CFM (top view). Multicolor images are generated by flipping the LED color, ensuring each frame is illuminated by a single color. The images are then superimposed and false-colored to create a composite image.

To demonstrate biological applications, we investigated immune cell adhesion with the cancer therapeutic drug Blinatumomab, a bispecific T-cell engager (BiTE)^30, 31^. Blinatumomab is a single-chain variable fragment of monoclonal antibodies containing a binding site recognizing CD3e (L2K-O7) and another site that recognizes CD19 (HD37)^30^. The BiTE molecule can trigger the activation of T-cells and localize those cells to CD19+ B-cells^32^. Blinatumomab is an FDA-approved drug to treat acute lymphoblastic leukemia^33, 34, 35^. By looking at BiTE-mediated binding between T-cells and CD19+ B-cells, we observed distinct time dynamics in the interaction profile that would be difficult to identify with other methods.

Our CFM-based approach provides the necessary statistics to quantitatively measure changes in cell adhesion across a wide range of conditions. Coupled with affordability and ease of use, our approach could broadly enable the study of critical cell processes, from fundamental cell biology to therapeutic cancer treatments like immunotherapy.

## 2. Methods

### 2.1 Materials

Nalm6 cells were purchased from ATCC (VA, USA). Jurkat E6-1 cells were a gift from Prof. Sizun Jiang. Poly-L-Lysine (PLL) coated coverslips (18 mm diameter, #1 thickness) were purchased from Neuvitro (H-18-PLL, WA, USA). Non-coated coverslips (18 mm diameter, #1 thickness) were purchased from VWR (48380-046, PA, USA). Blinatumomab recombinant antibody (MA5-41729) was purchased from Invitrogen (MA, USA). Soluble CD19 from Acro biosystem (CD9-H82F6, DE, USA) was used to block cell-bound CD19 from BiTE. The OKT3 anti-CD3 antibody was used to block CD3-BiTE interaction and shares 90% sequence homology with the BiTE aCD3 domain (Invitrogen 14-0037-82) ^36^. BSA-blocking solution was purchased from CANDOR Bioscience GmbH (BSA-block, Wangen, Germany). Jurkat and Nalm6 cells were grown in RPMI (ATC modification Thermofisher A1049101) supplemented with 10% FBS (ATCC 30-2020) and 100 U/ml Penicillin-Streptomycin (Thermofisher 15070063). Amyl acetate was purchased from Millipore Sigma (W504009-500G, MA, USA). Nitrocellulose was purchased from Biorad (#1620115). PBS buffer (10010-023, pH 7.4) was purchased from Thermo Fisher Science (MA, USA).

### 2.2 CFM Setup

The core setup of the centrifuge force microscope has been previously described^27, 37^, with updates for multicolor fluorescence provided in the updated parts list (**Supplemental Table 1**). A small microscope with an LED light source, sample holder, objective, and camera is assembled (**Figure 1B**). The microscope is held in a custom 3D printed holder, which fits into a commercial benchtop centrifuge (Heraeus X1R, Thermo Scientific). The centrifuge is modified with a fiber optic rotary joint and a computer control module, as specified in Yang et al., 2016^26^. The camera signal is sent out from the centrifuge through a fiber optic cable with a rotary joint, allowing free rotation. Rechargeable batteries power the entire setup. In contrast from the previously described setup, multiband pass excitation and emission filters were added along with a multicolored LED to modify the CFM for fluorescent imaging. A microcontroller (Trinket M0, Adafruit) was also added to sync the LED color with the camera to enable temporally multiplexed multicolor fluorescence imaging (**Supplemental Figure S1**). Trajectories are analyzed from 300 to 3000 rpm (13 - 1340 g) to avoid artifacts of shaking upon centrifuge start and nonlinear acceleration between 0 and 300 rpm.

The computer control module in the centrifuge allows the specification of ramp protocols by creating stair-step commands. The centrifuge is programmed to accelerate to a specific speed, and after it is reached, it is immediately instructed to increase to another speed. The size of the steps controls the ramp speed. In addition to the internal reporting of the centrifuge, the rpm was measured externally using a photodiode (OPB732, TT electronics, TX, USA) which measured the time between revolutions based on a piece of reflective material attached to the rotor. The force ramp protocol was implemented in the WinMass (ThermoFisher Scientific) centrifuge control software. Small incremental steps were used to achieve linear force ramps. The step size in the RPM controller script determined the loading rate.

### 2.3 Chamber preparation

The CFM sample chamber is made with double-sided Kapton tape (Kapton PPTDE-3) sandwiched between one coated (nitrocellulose or PLL) coverslip and one blank coverslip as a lid (**Figure 1C**). Double-sided tape is cut as an annulus, with a 7.5 mm inner diameter and 18 mm outer diameter. To increase the volume and depth of the sample chamber, three layers of tape are stacked on the coverslip to create a chamber with a volume of ∼20 μl. After the sample is prepared with cells, the lid is attached. The sample chamber is mounted on a supporting slide glass with Kapton tape to give structural support during preparation and centrifugation (SI Howard Glass Co, B-270 Ø 25 mm, 0.9 mm thick).

### 2.4 Staining of cells

Cells were grown to ∼1 million cells/ml in RPMI with 10% FBS and 50 U/ml Penicillin-Streptomycin at 37 degrees with 5% CO2. Cells were counted using the Luna II automated cell counter (Logos Biosystems). For staining, 10 ml of cells in media were centrifuged at 400 g for 2 minutes. The media was removed, and the cells were resuspended in 1 ml of PBS. To stain, 1 μl of CellTrace CFSE dye (Invitrogen C34554) or 2.5 μl of CellTrace far red dye (Invitrogen C34564) (prepared to manufacturer instructions in DMSO) were added. The cells were incubated at 37 degrees for 20 minutes. The cells were spun down again, the buffer was removed, and the cells were resuspended in 10 ml RPMI+FBS. The cells were incubated for at least 20 minutes prior to imaging.

### 2.6 BiTE molecule attachment on the nitrocellulose coverslip

A layer of nitrocellulose was deposited on the surface by thinly spreading a solution of 1% (w/v) nitrocellulose in amyl acetate on the coverslip surface (18 mm diameter, #1 thickness). The cover glass was incubated in the oven (65 °C) for 10 minutes.

For BiTE adsorption, a 30 μL droplet of PBS (pH 7.4) containing a specified BiTE concentration (0 to 125 nM) was added to the nitrocellulose coverslip surface. The chamber was incubated for 60 minutes at room temperature to allow adsorption, and the entire chamber was gently soaked in 4 ml of PBS buffer once to remove the free BiTE molecules from the BiTEs-attached sample surface. To minimize non-specific binding on the coverslip’s surface, the chambers were filled with 20 μl of buffer A (10 mM HEPES, pH 7.4, with 150 mM NaCl and 20% (v/v) BSA blocking solution) for 30 minutes at room temperature.

Jurkat T-cells were washed in buffer A and concentrated using centrifugation (400 g, 1 minute). Cells were resuspended in buffer A to 10 million cells/ml (counted using the automated cell counter (Luna II, Logos Biosystem, South Korea)). For measurements, 5 μl of Jurkat cells were added onto the BiTE-coated surface. The chamber was sealed with a lid coverslip on top.

### 2.6 Nalm6 monolayer attachment to the PLL cover glass

After staining, Nalm6 cells were concentrated to 40 million cells/ml in RPMI medium without FBS, and 30 μl were injected on the poly-L-lysine (PLL) coated coverslip surface. After 60 minutes of cell adsorption on the PLL coverslip, the excess cells were removed by flipping the chamber upside down in PBS for one minute. Fresh RPMI media with FBS was added to the sample chamber, and the sample chamber was incubated (37 C) for at least 30 minutes before starting use.

The chamber’s media was exchanged with 30 μl of buffer A mixed with the specified BiTE concentration. The cells were incubated (37C) for 5 minutes to allow BiTE binding to CD19 target antigens on the Nalm6 surface. After the incubation, 5 μl of buffer A + 10 million cells/ml Jurkat cells was added to the chamber. The chamber was sealed with a lid coverslip on top and incubated for the specified time before imaging. For blocking experiments, OKT3 was premixed with Jurkat cells for 5 minutes and OKT3 was also added to the monolayer at the same concentration before mixing. Jurkat cells for the cell-cell avidity experiment were prepared as described above.

### 2.7 Imaging

The images collected from the camera were recorded with custom LabView software. Images were taken at 4 frames/second with an exposure time of 0.2 seconds per frame. Image size was 4096×2160, and a 20x objective (with the casing removed) was used (170 × 170 nm/pixel).

Videos were analyzed using a custom processing pipeline that identified cells through a Regional-CNN based on the YOLO5 algorithm trained on a human-annotated data set^38, 39^. The total number of cells on each frame was counted. Small fluctuations in the number of cells detected frame to frame could occur due to slight shifting of the field-of-view and slight differences in cell-object detection from frame to frame.

## 3. Results

### 3.1 Fluorescence centrifuge force microscope: Instrumentation and cell assay overview

To enable high-throughput single-cell measurements, we developed a centrifuge force microscope (CFM) with multichannel fluorescence capabilities based on a core design previously introduced by our lab^26, 37^. Briefly, the CFM is a small microscope that fits into a 3D-printed holder that fits into a commercial centrifuge (**Figure 1B**). A fiber optic cable and slip-ring transmit the signal from the centrifuge for video recording during operation.

Multi-color fluorescence imaging was required to monitor multiple cell types during centrifugation. Single-channel fluorescent CFMs have previously been used to study colloids^40^. Here, we describe a novel CFM design with dual-channel fluorescence capabilities. A multi-color LED (red and blue) was set up in transillumination below the sample chamber (**Figure 1B** and **S1**). A multi-bandpass excitation filter was inserted after the LED to filter out non-excitation wavelengths. To optimize image quality, a small aperture was included on either side of the excitation filter to block non-orthogonal light rays. LED illumination excited the distinct dyes in each cell type (Jurkat cells and monolayer cells) before the light passed through the objective lens. A multiband-pass emission filter positioned behind the objective allowed only the fluorescent light emitted by the dyes to reach the CMOS camera, enabling detection of the cell positions. The LED color was controlled by a microcontroller triggered by the CFM camera to switch colors after every frame, allowing both channels to be captured in alternating frames (**Figure 1D**).

Cells were fluorescently labeled with CellTrace, a low cytotoxic dye that covalently binds to free amines. The monolayer channel was used for quality control, assessing the coverage of the monolayer and watching for bubbles that could come through the field of view. The target cell channel was analyzed by counting the cells in each frame—a custom image processing pipeline identified cells through an RCNN neural net based on the YOLO5 algorithm^38, 39^.

After placing the cells onto the target surface, the chamber was sealed and loaded into the CFM (**Figure 1C**). Simultaneously, a video recording was started and the CFM was flipped as the microscope (**Figure 2A, Supplemental Figure S3** and **supplemental video file**) was loaded into the centrifuge bucket. The flip changed the gravity from pushing the cells into the surface to pulling away from the surface, applying a low force of ∼0.3 pN per cell.

**Figure 2.**
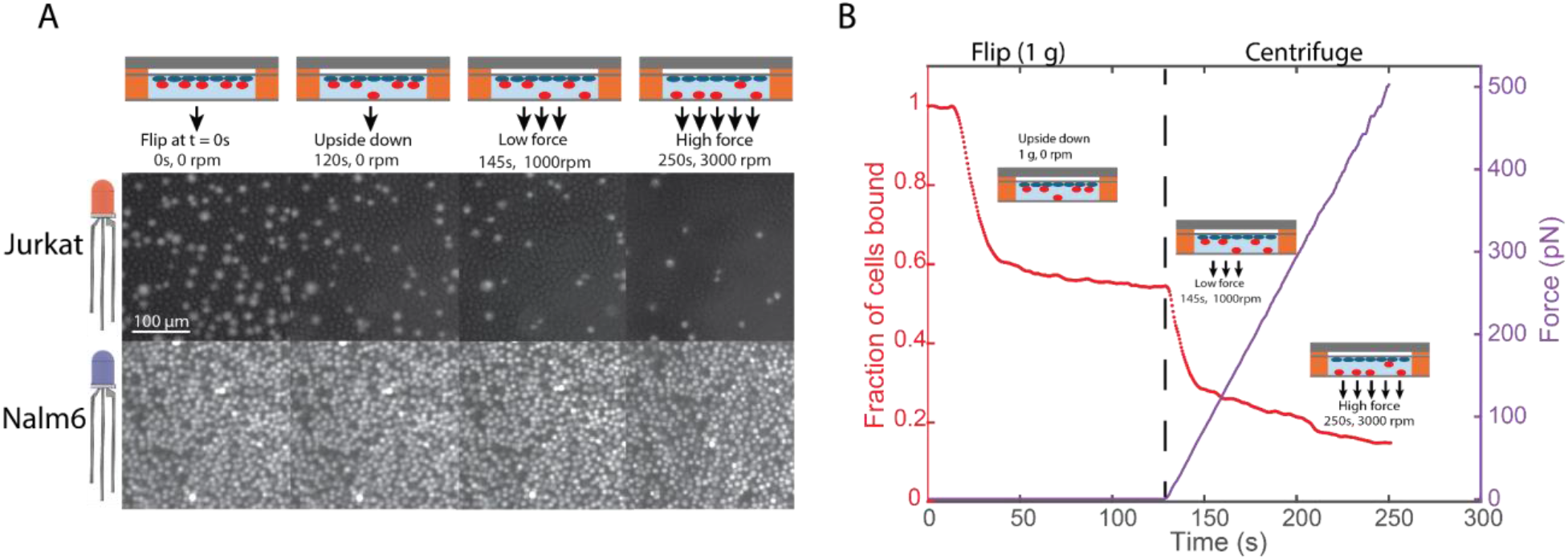
Fluorescence CFM detection of Jurkat T-cell detachment from a Nalm6 B-cell monolayer surface. **(A)** Partial field of view of Jurat T-cells (top) and Nalm6 B-cells (bottom) from the CFM at different time points during centrifugation. At t=0, the chamber was flipped upside down, allowing non-adherent cells to leave the surface. At t=120s, centrifugation began, and the force was ramped at a rate of 4 pN/s. As the force increased, Jurkat T-cells unbound and disappeared from the field of view. Example video included in supplemental material and **Supplemental Figure S3. (B)** Unbinding trajectory (red) of Jurkat T-cell detachment showing the fraction of cells bound, initially under the force of gravity then subsequently under a 4 pN/s force ramp. The applied force is shown in purple.

We generally observed an initial decrease in cell count during the flip, as non-adherent cells fell away due to gravity, before reaching a relatively steady level (**Figure 2B, Flip 1g section**). After two minutes, the centrifuge was activated and followed a linear force ramp protocol (**Figure 2B, centrifuge section**). An adhesion frequency was calculated as the ratio of cells remaining after the two-minute flip—prior to centrifuge activation—to the initial cell count. The adhesion frequency represents the fraction of cells exhibiting a minimum interaction strength, with the cells that detached due to gravity categorized as non-adherent.

The applied force on each cell is given by

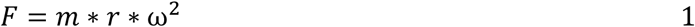

Here, F is the applied force, r is the distance from the rotation axis, ω is rotational speed, and m is the effective mass defined as m = *V*_*cell*_(*ρ*_*cell*_ − *ρ*_*water*_), where V is the volume of a single cell and *ρ* is density. The volume and density of the Jurkat cell line we studied are well-documented in the literature and have previously been approximated as spheres with a diameter of around 10 um and a density of 1.07 g/ml^41, 42, 43^. Using these parameters, we calculated the applied force on each cell at each measured rotational speed (**Figure 2B, purple line**). By controlling the rotational speed with WinMass (ThermoFisher Scientific), we obtained a linear force ramp between 2 to 16 pN/s, up to a maximum force of 500 pN per cell (**Supplemental Figure S2**).

### 3.2 Cell-protein measurements: Quantifying the strength of BiTE-immune cell interactions

Using the CFM method described above, we interrogated BiTE-immune cell interactions by attaching BiTE molecules to a surface, allowing cells to first bind before applying centrifugal force to measure their detachment lifetimes. By controlling protein identity and surface density, we initially validated the method to ensure reproducibility and characterize the specificity of binding interactions. We used the BiTE molecule Blinatumomab, which has one binding site for CD3e and another for CD19, enabling measurements of both Jurkat T-cells (CD3e+ domain of TCR) and Nalm6 cells (CD19+) (**Figure 3A**)^30, 44, 45^. Each BiTE molecule can bind to only one receptor of each type at any given time.

**Figure 3.**
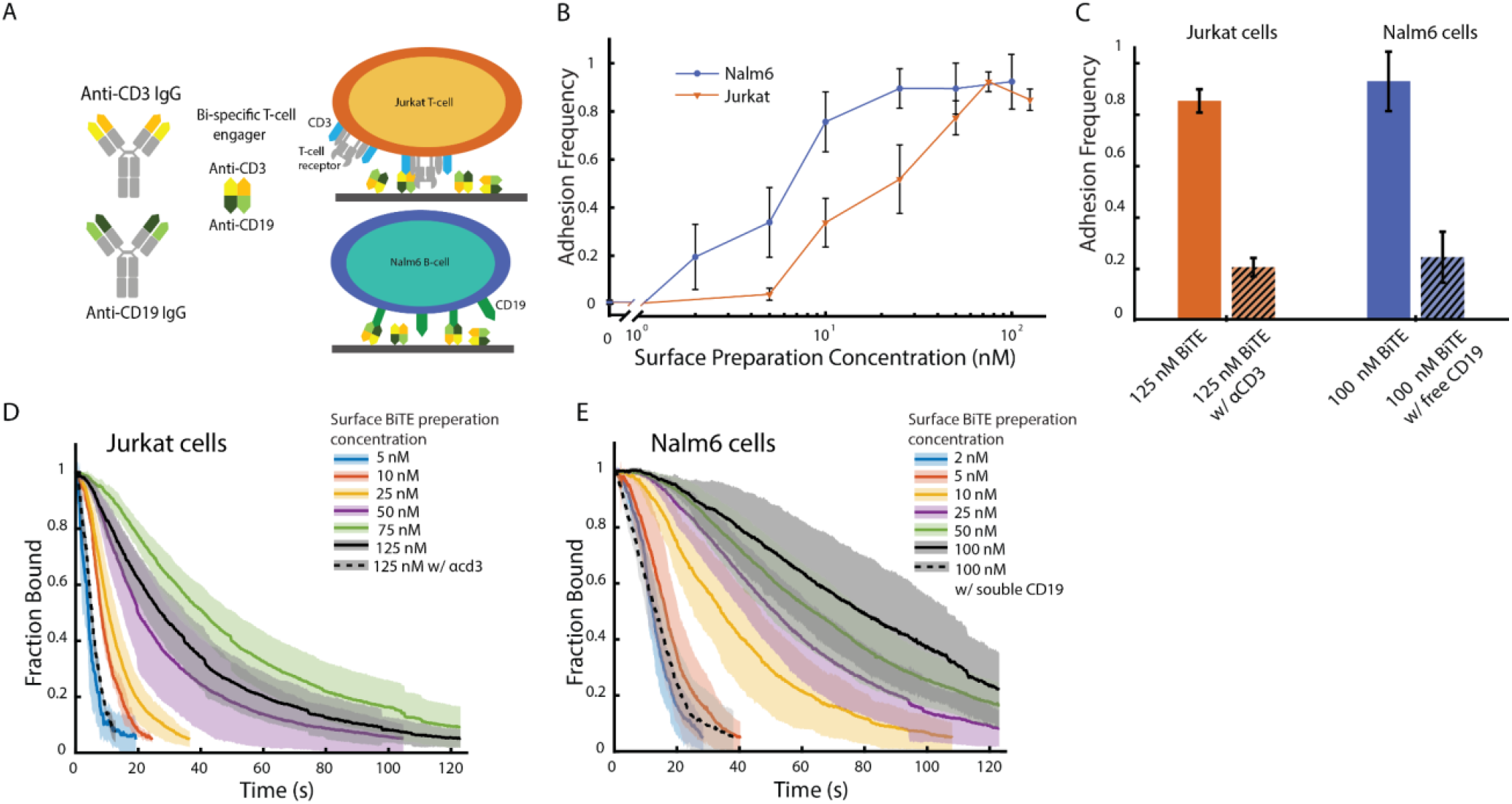
Jurkat T-cells and Nalm6 B-cell adhesion to BiTE functionalized surfaces. **(A)** Schematic image of cell attachment to the surface. BiTE (Blinatumomab) contains CD3ε and CD19 binding sites; it is randomly deposited on the surface and binds to Jurkat-T cells through the CD3 receptor and Nalm6 cells through CD19. **(B)** Jurkat (orange) and Nalm6 (blue) cell adhesion frequency vs. BiTE preparation concentration on the glass surface after 2 minutes of gravity. Adhesion frequency is calculated as the number of cells after the two-minute gravity flip divided by the starting number of cells. Given concentration represents the concentration of BiTE solution the surface was prepared with. Number of trials in order of increasing concentration (Jurkat, Nalm6): N_trials_ = [5, 5,3,5,5,5,5,3], N_trials_ = [3,4,5,3,5,5,4,3], Total cells observed: N_cells_ = [2582,3070, 3805,1818,2827,3389,2555,3186], N_cells_ = [3668,3202,1787,5045,3900,4458,4072,2826] **(C)** The blocking of Jurkat T-cell and Nalm6 cell bindings on BiTE functionalized surfaces from addition of anti-CD3 antibody and CD19 proteins, respectively. Number of trials: N_trials_ = [5,3,4,3] **(D)** Unbinding trajectories showing the fraction of Jurkat cells remaining bound on the surface with different BiTE concentrations in the presence of a 4 pN/s ramping force. Given concentration represents the concentration of BiTE solution the surface was prepared with. For each concentration, at least three runs were averaged, with the standard deviation shown as the shaded region. Number of trials: N_trials_= [5,3,5,5,5,5,3], Total cells observed: N_cells_ = [128,576,2595,3005,4150,3492,584]. Fewer cells survive the 2-minute 1x g flip at lower concentrations, so fewer rupture events were observed. Full force trajectories shown in **Supplemental Figure S4. (E)** Unbinding trajectories showing the fraction of Nalm6 cells remaining bound on the surface with different BiTE concentrations under 4 pN/s ramping force. Number of trials: N_trials_ = [4,5,3,5,5,4,3], Total cells observed: N_cells_ = [622, 1263, 1406, 2518, 3023, 2369,799]. Full trajectories are shown in **Supplemental Figure S5**.

We saw a strong relationship between the concentration of BiTEs in the surface preparation and the resulting adhesion frequency (**Figure 3B**). When no BiTE was added, more than 99% of the cells fell off the surface during the initial flip, indicating effective passivation with BSA blocking solution. For both cell types, increasing the BiTE concentration resulted in a higher adhesion frequency. At high concentrations, adhesion frequency plateaued, with more than 90% of the cells remaining after the initial flip. Adding soluble anti-CD3 antibody (OKT3) or soluble CD19 to the buffer competed with the BiTE-receptor interaction on their respective target cells (**Figure 3C**). This competition resulted in a significant decrease in cell adhesion for both Jurkat and Nalm6 cells (from ∼80-90% to ∼20%), supporting that the BiTE-receptor interaction drove the observed adhesion.

After the initial flipping of the CFM, the centrifuge was spun to apply a linear force ramp of approximately 4 pN/s at time T = 120s (**Figure 2B**, purple line). **Figure 3D-E** illustrates the number of Jurkat or Nalm6 cells remaining as a function of time, normalized by the number of cells on the surface at the start of centrifuge acceleration. For each condition, multiple trials are averaged at each time point, with the standard deviation shown as a shaded band. Full trajectories of each ramping experiment are available in **Supplemental Figures S4** (Jurkat cells) and **S5** (Nalm6 cells).

As the BiTE concentration increased, the lifetime of both cell types under force also increased, with cells requiring higher forces to be detached from the functionalized surface. Nalm6 cells exhibited longer lifetimes under force at a given BiTE concentration, possibly due to a greater number of receptors or to higher per-receptor affinity. For the Jurkat cells, fraction-bound trajectories at preparation concentrations above 75 nM did not show increased binding strength, suggesting that the available binding sites were saturated. Saturation was observed in both the adhesion frequency and force ramping data, with a 125 nM BiTE concentration yielding a similar response to 75 nM. Blocking with aCD3e or soluble CD19 resulted in weaker adhesion, with the population rupturing at low forces (**Figure 3D-E**, dotted lines).

Each field of view contained approximately 500-1000 cells, and each trial took roughly 15 minutes. This high-throughput approach enabled us to capture over 25,000 single-cell unbinding events, allowing the construction of detailed rupture event distributions (**Supplemental Figure S6**). The method’s statistical power enables robust quantitative comparisons of binding behaviors across various conditions and cell types.

### 3.3 Kinetic Analysis of Cell Binding

To further investigate the dynamic strength of cell-BiTE interactions, we analyzed the kinetics of cell adhesion under increasing BiTE concentrations. At higher surface-BiTE concentrations, we observed both an increase in the number of cells that remained after flipping under gravity, and longer lifetimes under force of the cells that remained. Since both adhesion frequency and binding lifetime arise from molecular interactions such as the number of receptor-ligand bonds per cell, we analyzed their relationship to determine how they are governed by these shared molecular properties.

We found that the fraction of remaining cells, f(t), under a linear force ramp (**Supplemental Figure S7**), was well-described by a stretched exponential function of the form,

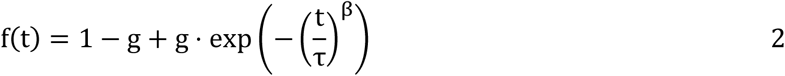

A stretched exponential is a generalization of regular exponential decay that allows for more flexibility in modeling failure processes where the decay rate changes over time ^46^. In this case, the time constant **τ** provides a single metric that characterizes the population’s lifetime under specific ramping conditions, similar to how it would in a regular exponential decay. The β factor accounts for changes in the off-rate over time—here, the continuously increasing force accelerates the off-rate, effectively compressing the decay relative to a standard exponential. Finally, the offset term g represents the fraction of cells that remain bound to the surface.

Plotting the adhesion frequency against the fitted population lifetime **τ** for each trajectory reveals a universal curve for a given force ramp and receptor type (**Figure 4A-B**). This relationship holds true regardless of surface concentration or variability between individual surface preparations, demonstrating a strong correlation between adhesion frequency and cell binding lifetime. Interestingly, blocking the binding site with a competitor reduced both the adhesion frequency and lifetime but preserved the relationship between them. The strong correlation suggests that a shared factor, likely the number of available receptors, governs binding.

**Figure 4.**
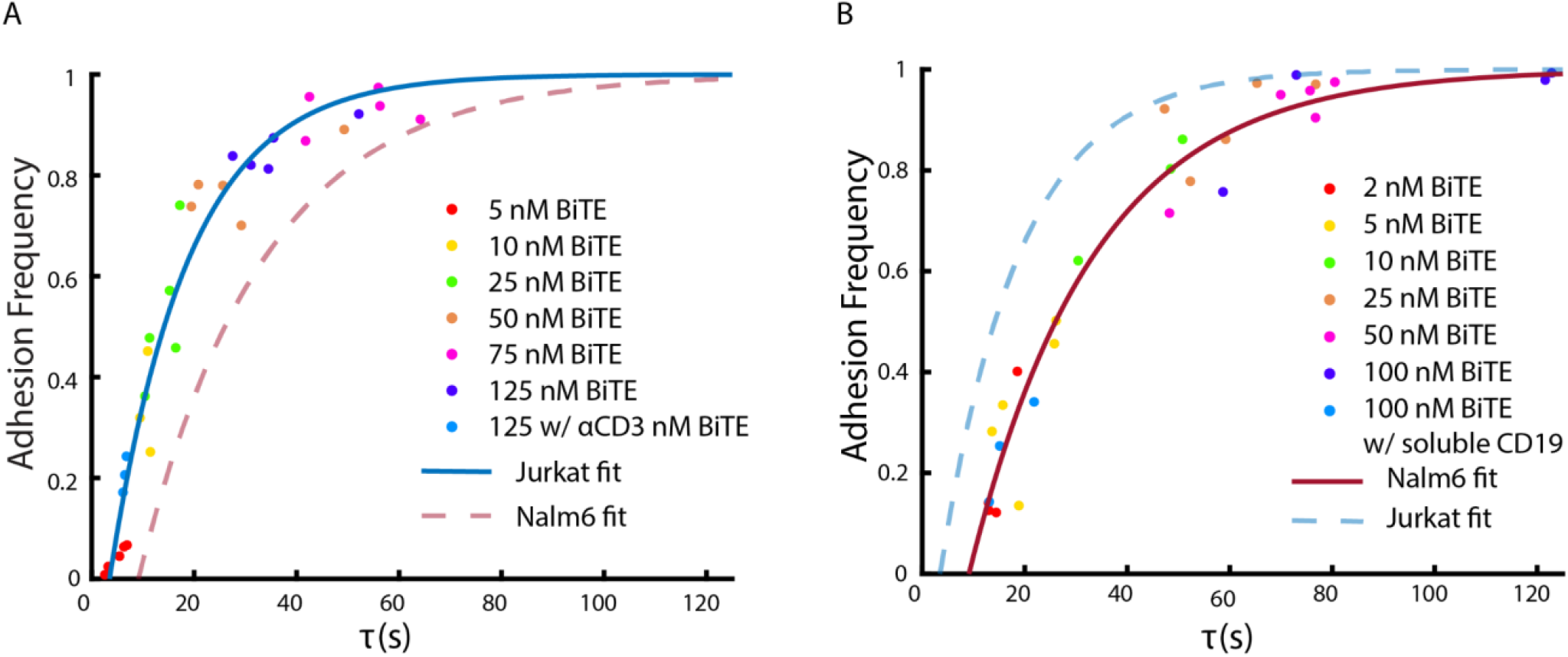
Binding analysis of surface functionalized with Blinatumomab BiTE to different cell types. **(A)** Each dot represents a single trial of Jurkat cells binding to BiTE, with surface concentration indicated by color. Adhesion frequency is calculated as the fraction of cells remaining bound after gravity relative to the initial number. The unbinding trajectories of cells under force (**Figure 3C-D, Supplemental Figure S4-5**) are fitted to a stretched exponential function: 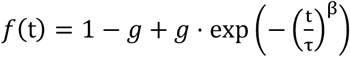. The fitted lifetime parameter (**τ**) is plotted against adhesion frequency for each trial. To fit a universal curve to the data, a set of parametric equations based on the distribution of bonds per cell (given by λ) is used (see text). The predicted adhesion frequency is modeled as AF(λ)=1−CDF.Poisson(0,λ) and the predicted population lifetime is modeled by τ(λ) = *k*λ + *x*_0_, with k and *x*_0_ as free parameters. The fitted curve (solid line) has parameters k=15.4 *x*_0_=3.5 (RMSE: 0.07). The dotted line represents the analogous curve from the Nalm6 binding data in B for comparison. Number of trials: N_trials_ = 31, Total cells observed: N_cells_ = 23,232. **(B)** Nalm6 cells binding to a BiTE-functionalized surface, with data processed as explained in **4A** showing the relationship between fitted lifetime **τ** and adhesion frequency. The solid line shows the fit of the Nalm6 data (k = 24.5, *x*_0_ = 9.1, RMSE: 0.08), and the dotted line shows the Jurkat fit. Number of trials: N_trials_= 29, Total cells observed: N_cells_ = 28,958

Intuitively, having more receptors increases the likelihood of cells adhering and enhances cell binding strength by providing more opportunities to form bonds.

To model the relationship, we assume that the number of bonds formed on each cell follows a Poisson distribution characterized by the expected rate of bond occurrence, λ, and the number of bond occurrences, x,

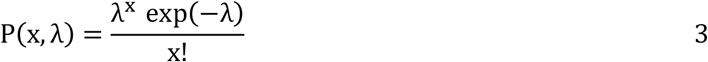

The adhesion frequency represents the probability that at least one bond forms, ensuring survival during flipping. This fraction is calculated using the complement of the cumulative distribution function (CDF) of the Poisson distribution evaluated at zero (x = 0), expressed as AF(λ) = 1 − CDF_poission_(0, λ).

During the force-ramp phase, the fitted population lifetime **τ** of the cells increases as more bonds are added. For independent, non-cooperative bonds loaded in parallel under a linearly ramping force, the lifetime of each cell scales approximately linearly with the number of bonds, provided the bond count is sufficiently large^47, 48^. If the number of bonds is Poisson distributed, the fitted population lifetime **τ** is approximately proportional to λ, scaled by a proportionality constant k. Additionally, an offset *x*_0_ is included to account for the lifetime not being zero when the adhesion frequency is zero. Thus, we can approximate the population lifetime by the expression **τ**(λ) = k * λ + *x*_0_, where k and *x*_0_ are free parameters. The parameter k characterizes the strength of individual bonds, with a higher k value indicating a slower off-rate under these force-ramping conditions (**Supplemental Figure S8**).

Given the relationships for population lifetime (**τ**) and adhesion frequency (AF), we have established a set of parametric equations that relate adhesion frequency and population lifetime. By varying the surface concentration, we effectively sample different values of λ. These relationships were used to fit the experimental data, with k and *x*_0_ as the two free parameters representing aspects of individual receptor lifetimes and receptor densities. The fitted curves for Jurkat and Nalm6 cells were overlaid on the respective datasets (**Figure 4A-B**).

These universal curves differ between the two cell types, reflecting their dependence on the specific receptor and capturing the distinct avidity of each cell type. The fits suggest that the CD19-BiTE interaction is stronger than the CD3e-BiTE interaction (CD19:k=24.5, CD3e:k=15.4). Intuitively, Nalm6 cells (CD19+) survive longer at a given adhesion frequency than Jurkat cells (CD3e). The trend is supported by the literature, which gives dissociation constant (Kd) values for 1.49 × 10^-9 M (Blinatumomab BiTE vs. Nalm6) and 2.6 × 10^-7 M (Blinatumomab BiTE vs. purified human T-cell) ^30^. When measuring only one receptor type at a time, the model enables comparison of relative binding strengths between different receptors without requiring explicit measurement of receptor density on the cells.

We also demonstrated the ability to explore different force loading rates with the CFM (**Supplemental Figure S9)**. Ramp protocols were created to apply various loading rates between 2 pN/s and 16 pN/s. The Jurkat T-cell BiTE interaction was measured over a range of loading rates at 10 nM BiTE, balancing statistics and low bond/cell number. The off-rate at a given force was calculated based on the number of cells that detached at a given force, normalized by the number of cells available and scaled by the time interval^49, 50^. As expected, the off-rate increased as the force increased. However, the observed off-rate plateaued as the number of cells decreased, possibly due to an increase in the fraction of multiply-bonded cells. The technique demonstrates the CFM’s capability for measuring cell unbinding kinetics at different forces and loading rates.

### 3.4 Cell-cell Measurements: Jurkat cell adhesion on Nalm6-BiTE monolayer surface

Next, we used the CFM to interrogate BiTE-mediated cell adhesion of Jurkat T-cells to Nalm6 B-cells, measuring the interactions of immune cells at the single-cell level. Nalm6 B-cells were attached to a glass cover slip to form a dense monolayer, then incubated with BiTE at the specified concentration; T-cells were then added on top. The cells were initially flipped to obtain the adhesion frequency based on the fraction of remaining cells after the two-minute flip, after which the force was increased at a loading rate of 4 pN/s as previously described. Based on the results in **Sections 3.2-3.3**, which provide evidence that both cell types bind to a specific site of the BiTE molecule **(Figure 3B-E**), we hypothesized that the BiTE would act as a bridge between the two cell types.

Surprisingly, changes in BiTE concentration had little observable effect on adhesion frequency within the tested range (**Supplemental Figure S12A**). Additionally, unlike the cell-surface measurements, the cell-cell adhesion measurements exhibited a relatively high background binding rate of approximately 30% even with BSA blocking solution. This may have been due to the presence of various proteins, lipids, and sugars on the surface of the monolayer cells, interacting specifically or non-specifically with the Jurkat cells. High background levels and trial variability may have obscured the relationship between BiTE concentration and adhesion frequency.

In order to determine the impact of BiTE concentration on adhesion strength, we examined the lifetime of each cell under a linear force ramp of approximately 4 pN/s. From the resulting survival fraction trajectories (**Figure 5A**), we observed that increases in BiTE concentration up to 75 nM enhanced the adhesion strength between the two cell types. As a negative control, we pre-incubated Jurkat T cells with an anti-CD3 antibody to block the BiTE-CD3 interaction, which reduced adhesion back to the 0 nM control level (**Figure 5A**, back dashed curve). This data supported the key role that BiTE bridging played in driving the increase in adhesion strength.

**Figure 5.**
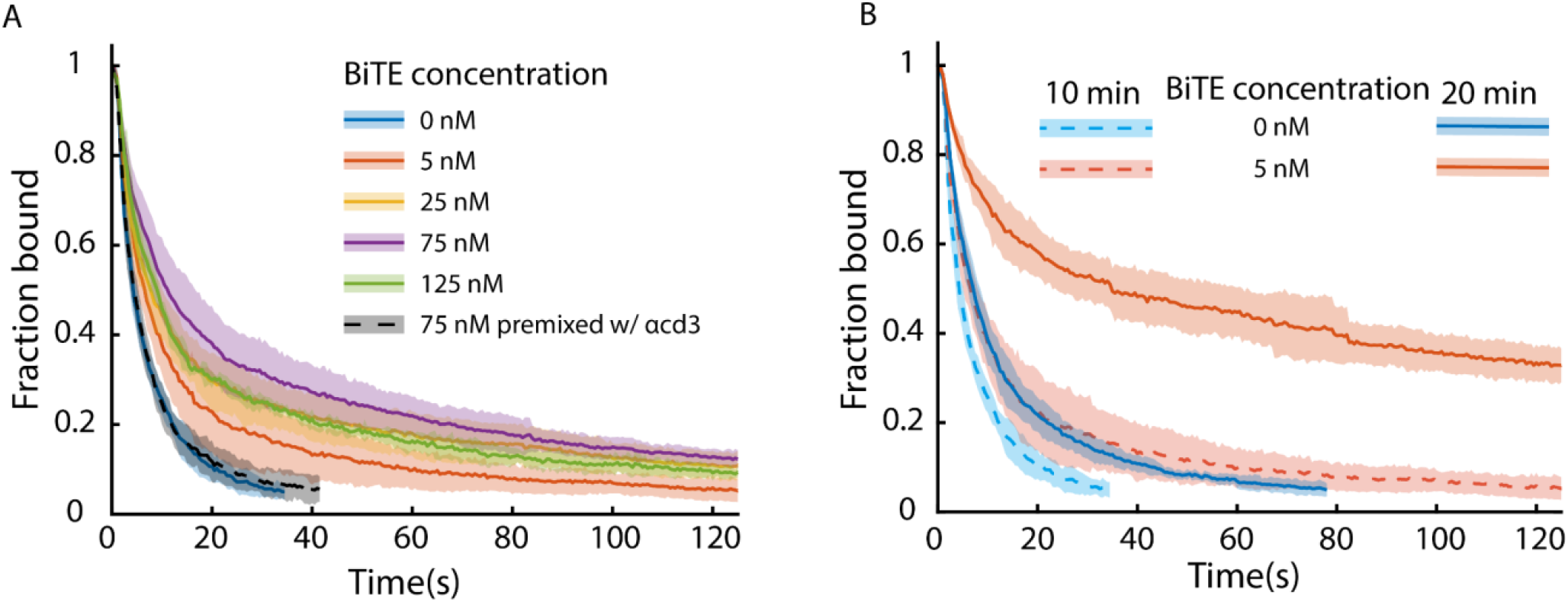
Jurkat T-cell strength on BiTE treated Nalm6 cell monolayer. **(A)** Unbinding trajectories of Jurkat T-cells bound to a Nalm6 monolayer under 4 pN/s ramping force at different BiTE concentrations. Jurkat cells were incubated with 10 minute attachment times on the monolayer for all concentrations. Given concentration represents the concentration of BiTE in buffer mixed with monolayer. For each concentration, at least three runs were averaged together, with the standard deviation presented as the shaded region. Number of trials: N_trials_ = [6,4,3,5,3,4], Total cells observed: N_cells_ = [1946,1578, 1447, 1909, 1036, 1138]. Full trajectories under force in **Supplemental Figure S10. (B)** Unbinding trajectories of Jurkat cells bound on Nalm6 monolayer at 10 and 20 minute attachment times and at two different BiTE concentrations (0 nM, 5nM) with 4 pN/s force ramp. Data listed: [0 nM 10 min, 5 nM 10 min, 0 nM 20 min, 5 nM 20 min]. Number of trials: N_trials_ = [6,4,4,5], Total cells observed: N_cells_ = [1946,1578, 1946, 2650]. Full trajectories under force in **Supplemental Figure S11**.

Leveraging the CFM’s ability to assay cell binding in parallel, we investigated whether BiTE binding influences cell adhesion over time. We applied force to capture potential adhesion dynamics by measuring binding at discrete time points along a cell’s trajectory. When the attachment duration was extended from 10 to 20 minutes, it resulted in a marked increase in cell adhesion. Under a linear force ramp, a much greater fraction of cells remained bound at high forces following 20 minutes of BiTEs incubation compared to 10 minutes. However, at the 20 minutes attachment time, adhesion showed little concentration dependence across the tested BiTE range (**Supplemental Figure 12B**). Notably, 5 nM BiTE at 20 minutes led to stronger adhesion than 75 nM BiTE at 10 minutes, underscoring the significant effect of attachment time. Despite the increased strength, increasing attachment time from 10 to 20 minutes caused minimal changes to the adhesion frequency (**Supplementa**l **Figure 12B**).

To explore whether longer attachment times enhance adhesion strength through a cellular mechanism, we tested incubation with the activating anti-CD3 antibody OKT3 (**Supplemental Figure 13B**) ^51^. Since OKT3 binds exclusively to CD3 on Jurkat cells, it should not mediate direct adhesion to Nalm6 cells. As previously shown, OKT3 can block Jurkat-Nalm6 adhesion mediated by BiTE binding with 10 minute attachment times (**Figure 5A**, black dashed curve). Interestingly, Jurkat cells incubated with only OKT3 for 20 minutes exhibited adhesion levels above the negative control, suggesting that some of the increased adhesion at 20 minutes in the BiTE experiments arises from T-cell activation rather than direct bridging alone. These results highlight the ability of the CFM to measure adhesion dynamics under different conditions to illuminate mechanistic differences underlying variations in T-cell binding.

## 4. Discussion

### 4.1 BiTE-mediated T-cell B-cell avidity

Comparing BiTE binding in cell-cell versus cell-surface experiments reveals distinct differences in adhesion responses at different stages throughout the force ramp. We observed a lower overall adhesion frequency in the cell-cell case but a higher proportion of cells surviving at high forces (**Figure 3C-E**,**5B, Supplemental Figure S12**). In surface-functionalized experiments, cell binding was generally homogeneous, with higher adhesion frequencies correlating well with adhesion strength under force (**Figure 4**). In contrast, in cell-cell experiments, a significant fraction of cells detached immediately after flipping; however, a significant percentage of cells that remained post-flipping withstood the highest applied forces (**Figure 5**). The coexistence of a large population with weak adhesion and a smaller subset with much stronger adhesion suggests the presence of a distinct subpopulation with a different response to BiTE in the presence of Nalm6 B cells. Further analysis using transcriptomic or proteomic profiling could help explain these cell-state differences.

Measuring cell-cell interactions between Jurkat and Nalm6 cells reveals minimal BiTE dependence on adhesion frequency. While surface functionalization can artificially increase receptor concentration, natural receptor abundance ultimately constrains cell-cell interactions. Consequently, beyond a certain threshold, increasing BiTE concentration may have little effect on adhesion, as the interaction is limited by the intrinsic densities of CD19 and CD3 receptors. Given the relatively high background signal for the cell-cell binding assay, adhesion frequency alone struggles to distinguish binding differences in the absence of force perturbation, highlighting the necessity of the CFM force ramp protocol.

The strong time dependence of adhesion raises the question of what primary mechanism drives the sudden increase in adhesion strength observed in a subset of the population. Jurkat cells binding to a BiTE-coated glass surface exhibited significantly lower retention under force than Nalm6-Jurkat at 20 minutes, with only 9 ± 7% remaining at 500 pN, even under saturated adhesion conditions (**Figures 3D, 5B**). This suggests that the increased avidity of Jurkat T cells on Nalm6 cells over time cannot be fully explained by BiTE-CD3 binding alone. Additionally, no time dependence was observed in the cell-surface assay (**Supplemental Figure S14**), suggesting that BiTE-CD3 binding reaches equilibrium within 10 minutes.

One possible mechanism for the increased avidity with longer attachment times is the lateral diffusion of CD3e and/or CD19 within the plasma membranes of Jurkat T-cells and Nalm6 cells, allowing for enhanced BiTE-mediated bridging. Previous findings show that TCR activation can induce the formation of TCR microcluster (TCR-MCs, 50-300 TCR molecules/cluster) ^52^. These microclusters are known to be laterally mobile in the membrane, enabling the recruitment of additional receptors to the interface^52, 53^.

Another potential mechanism involves CD3z-dependent activation of Jurkat T-cells, which can enhance binding by recruiting additional protein binders or reorganizing existing interactions. Blinatumomab, a known T-cell activator, initiates signaling pathways that lead to the formation of an immunological synapse^32^. The formation of this immune synapse is regulated not only by the lateral movement of TCR, LFA-1, and other membrane proteins but also by actin polymerization within the cytoskeletal network^53, 54^. Previous studies have shown that T-cell activation can induce actin-dependent centripetal movement of TCR and integrin microclusters, resulting in an immune synapse at the adhesion site^11, 53^.

Supporting the activation mechanism, we observed a slight increase in avidity after a 20 minute attachment period when Jurkat cells were incubated with the anti-CD3 activating antibody (OKT3) (**Supplemental Figure S13B**). Since OKT3 binds specifically to CD3+ T cells, any increase in binding is likely due to a cellular response rather than direct bridging. Notably, previous studies have reported a similar time-dependent increase in T-cell adhesiveness upon incubation with OKT3, consistent with our findings^55, 56^.

Our findings using the Centrifuge Force Microscope indicate that the strong cell adhesion observed reflects a dynamic avidity response in Jurkat T-cells interacting with Nalm6-BiTE cells. Notably, we observed significant changes in binding strength at low BiTE concentrations, aligning with clinically relevant concentrations used in patient treatments^30, 57^. Furthermore, the pronounced binding differences at longer attachment times became apparent only under force, underscoring the importance of measuring cell populations under force at specific time points to capture these critical dynamics.

### 4.2 Cell avidity force measurement tools

Built on a commercial microscope, the CFM enhances previous centrifuge-based techniques by providing improved temporal and force resolution for cell adhesion measurements. Centrifuge-based cell-cell adhesion assays have been used to quantify adherent cells before and after centrifugation, providing bulk measurements at a constant force^58, 59, 60, 61, 62, 63, 64^. However, these approaches, without live imaging of cell detachments, require prior knowledge about the relevant force range, do not capture the full population unbinding trajectories, and only provides population-level data.

The fluorescence CFM overcomes these limitations by integrating real-time imaging to capture complete cell detachment trajectories and employing a dynamic force ramp that eliminates the need for prior knowledge of the relevant force range (**Supplemental Table 2**). The current CFM design operates at speeds up to 3,000 rpm (∼1,600 g), generating forces up to ∼500 pN for 10 μm cells while enabling parallel measurement of over 1,000 cells. A force ramp (1–16 pN/s) rapidly probes a broad force range to determine the relevant force thresholds for different interactions. The CFM can also perform constant force experiments by employing a fixed rotational speed, facilitating measurements at physiological force levels for over 30 minutes.

Real-time monitoring yields high-quality data with continuous quality control, such as tracking dynamic cell detachment from monolayer cells or functionalized surfaces (**Figure 2A**). The camera ensures proper monolayer adherence and detects disruptions, such as bubbles affecting cell-surface attachments. By enabling single-cell tracking, the CFM shifts experiments from bulk assays to single-cell analysis, capturing the entire distribution of cell lifetimes and identifying distinct cell populations. Incorporating multicolor imaging in the fluorescence CFM allows multiple cell types to be measured simultaneously or dyes to be included to monitor other processes, such as ion uptake.

Cell force spectroscopy assays provide a critical complement to existing methods for studying cellular interactions. Techniques like fluorescence imaging and chemical tagging provide detailed information about cell types, receptor identities, and receptor organization. However, standard fluorescence imaging approaches are typically limited to identifying binding versus non-binding events rather than quantifying interaction strength^29^. Similarly, while conventional flow cytometry is commonly used to measure equilibrium binding between proteins and cells, it generally lacks the temporal resolution to capture binding kinetics^65^. In contrast, force-based cell adhesion assays are specifically designed to quantify binding strength between cells.

Our approach also complements other recently developed parallel cell avidity measurement techniques, such as acoustic force spectroscopy (AFS) and hydrodynamic flow systems, which face several challenges including (1) non-uniform force fields, leading to spatial heterogeneity and unpredictable force vectors; (2) reliance on external calibration for force quantification and (3) high instrument and operational costs, which can limit accessibility^18, 23, 24, 66, 67^. Centrifuge force microscopy, however, addresses these limitations and offers several advantages.

First, the centrifuge applies a uniform force field across the surface, with a straightforward relation between rpm and applied force (**Eq. 1**), virtually eliminating spatial differences in the direction and magnitude of force application across the sample. Force quantification is crucial in cell-cell measurements, where surface height and material properties could affect the force experienced by target cells under flow or acoustic waves. Unlike AFS, which relies on relative force calibration with standards like 10 μm polystyrene beads ^68^, centrifugation provides direct force quantification. Additionally, the CFM is a highly cost-effective and accessible solution, using standard lab equipment to enable the creation of a basic system for under $1,000--this reduces the cost barrier compared to most commercial optical or magnetic tweezers, AFM, or AFS systems, which can exceed $100,000^26, 69, 70, 71^. The CFM assay uses inexpensive, disposable cover slips, avoiding costly reusable chambers. This design supports rapid prototyping, diverse surface chemistries, and easier cleanup compared to AFS flow-through chambers. The protocol is straightforward, with centrifugation as the primary step, reducing the need for specialized expertise.

Further enhancements to the CFM could include increasing force ceilings and reducing variability from cell size and density differences. Faster centrifuges could achieve higher forces but would require imaging components that withstand high-g accelerations. Enhancing the density contrast between cells and medium could amplify forces but would necessitate careful osmotic regulation to prevent cell damage. Incorporating single-cell volume estimation could reduce force quantification error^41, 42, 43^. Enhancing imaging parameters such as resolution, illumination strength, and field-of-view stability would further improve data quality and expand analytical capabilities.

### 4.3 Future uses of CFM cell avidity measurements

The CFM is well-suited for studying mechanical forces in cellular processes, including lymphocyte activation pathways^72^. Parallel measurements complement optical trap studies by enabling broader experimental condition testing^73^. Fluorescence imaging allows real-time monitoring of T-cell activation under physiological forces. Measuring adhesion strength under these conditions could provide insights into how initial weak forces can influence long-term avidity. Beyond force ramp experiments, the CFM can also perform long-duration force clamps (30+ minutes) to study lifetimes at physiological forces and catch bond behavior^74^.

Even with the complexity of cell-cell interactions, comparing rupture curves across different conditions could provide mechanistic insights into cell avidity. Assessing relative binding strength allows for differentiation between cell responses or types, such as identifying adhesion-related proteins in knockout libraries. Current CFM methods could reasonably screen ∼100 knockouts, correlating interaction strength with knocked-out proteins to dissect receptor contributions and uncover underlying mechanisms.

The ease and statistical power of CFM assays make them valuable for clinical applications. The CFM could assess immune cell binding in patient samples, correlating binding strength with disease state, age, genetics, transcriptomics, or drug response. Selecting cells based on specific binding strengths could aid immunotherapy development, particularly in cancer, where moderate receptor binding affinities are often optimal^75^. The CFM’s precise force quantification enables targeted screening of desired binding strengths, making it a valuable tool for optimizing Chimeric Antigen Receptor (CAR) T-cells. By measuring and selecting CAR T-cells based on their mechanical binding profiles, the CFM could help improve efficacy and enhance therapeutic outcomes.

## 5. Conclusion

Our work introduces a high-throughput centrifuge force microscope (CFM) assay for studying cell-cell interactions at the single-cell level, featuring multichannel fluorescence for expanded imaging capabilities. This method enabled the capture of time-dependent and BiTE-specific effects on Jurkat-Nalm6 interactions–insights that are difficult to obtain with other techniques. With its accessible design and ability to perform hundreds to thousands of force measurements in parallel, the CFM is well-suited for investigating adhesion mechanics under force, offering valuable insights into cancer research and immunology.

## Data Availability

The datasets generated during and/or analyzed during the current study are available from the corresponding author on reasonable request.

## Code Availability

Code is available upon reasonable request.

## Acknowledgments

W.P.W. thanks NIH NIGMS (R35GM119537). We thank Dr. Andrew Ward for assistance with Labview and Prof. Sizun Jiang for Jurkat cells. K.K. thanks Dr. Marijn van Loenhout and Dr. Serena Davoli for helpful advice. We thank Dr. Ken Halvorsen and Prof. Xin Zhou for helpful discussions and suggestions.

## Author contributions

H.T.B, K.K and W.P.W were involved in idea conception, experimental design, data analysis, collaborative discussions, and manuscript writing. H.T.B and K.K carried out experiments.

