## Supplemental info for "High-Throughput Centrifuge Force Microscopy Reveals Dynamic Immune-Cell Avidity at the Single-Cell Level"

### Supplemental Figures S1-S14, Tables 1 and 2:

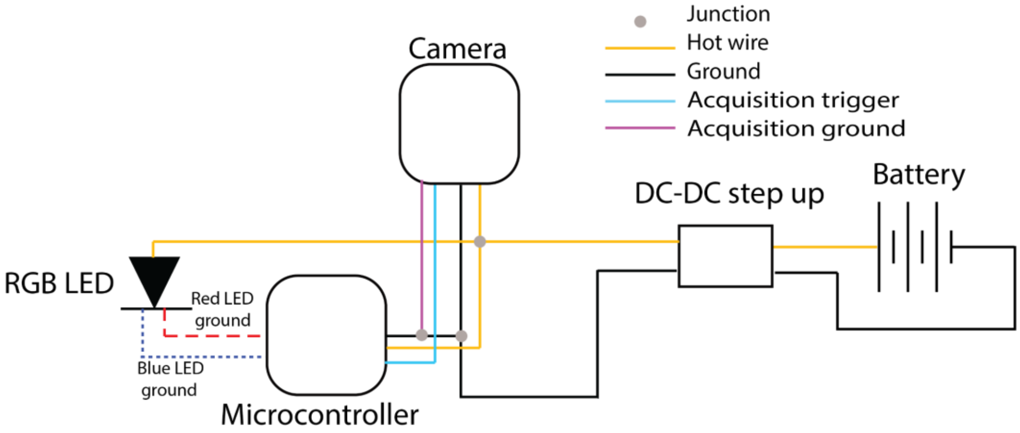

**Figure S1**. Circuit diagram for multicolor LED and microcontroller. Color is programmed by the microcontroller, which switches color based on the camera frame trigger. A detailed list of parts can be found in the **Supplemental table 1**. Similar to previous CFM designs, a 5V DC step-up converter is connected to the battery to maintain a constant voltage supply. This setup powers the camera, LED, and microcontroller. The camera's Pin 4 (opto-isolated output) is wired to an interrupt pin on the microcontroller, which detects a rising signal. The camera is configured to send a signal at the beginning of each frame. The LED's color is determined by grounding a specific colored lead, allowing current to flow and control the LED color. Each colored LED lead is connected to a pin on the microcontroller, which determines whether current flows and the resulting color. This configuration enables the LED color to sync with the camera's frame acquisition, illuminating only one color per frame. For protein-cell experiments, only one fluorescence channel was used.

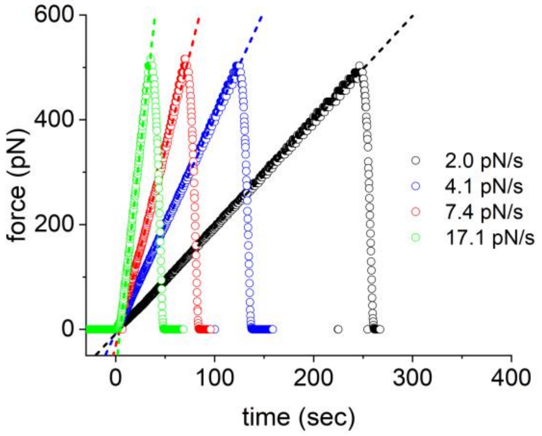

**Figure S2**. Four different linear loading rates of CFM by controlling the operation program. Loading rate calculation based on a spherical cell of diameter 10 microns, a buffer density of 1.00 g/ml, an estimated cell density of 1.07 g/ml, and a distance from the rotation axis of 15 cm.

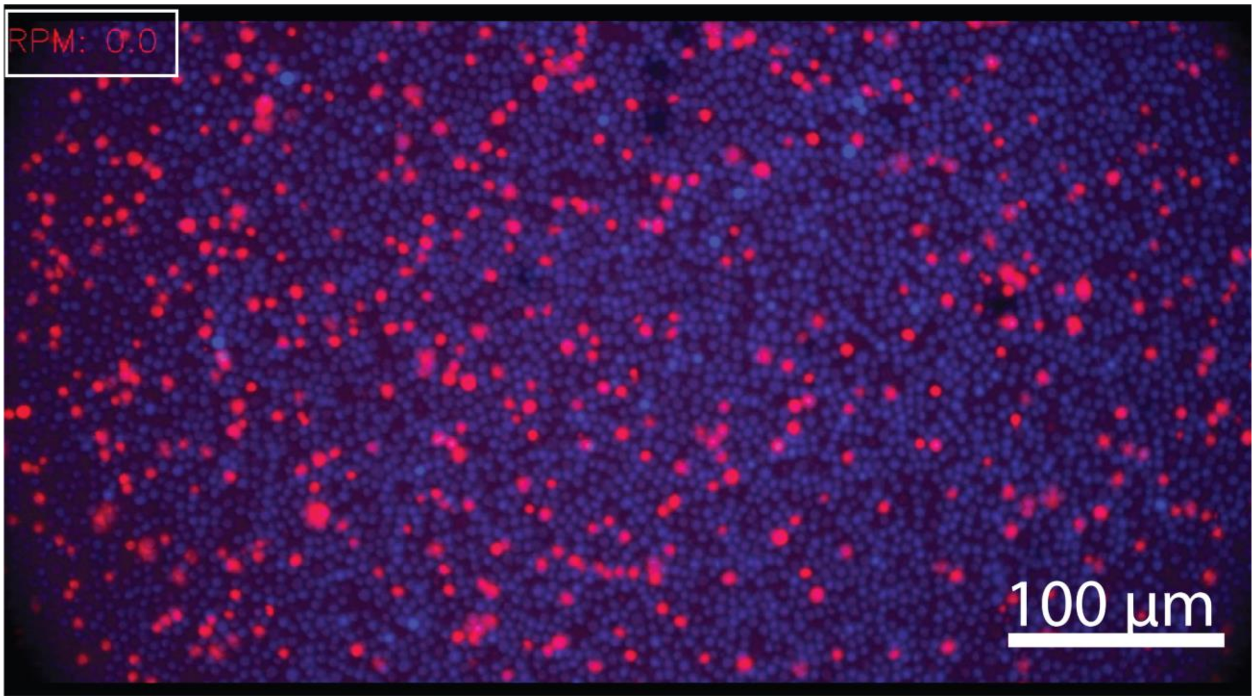

**Figure S3**. Composite image of Nalm6 cells in a dense monolayer (blue) with Jurkat cells on top (red). Full video of example included as supplemental file. RPM in top right corner shows real time RPM of centrifuge at time the image was taken. Video file is sped up 13x and images compressed to reduce file size.

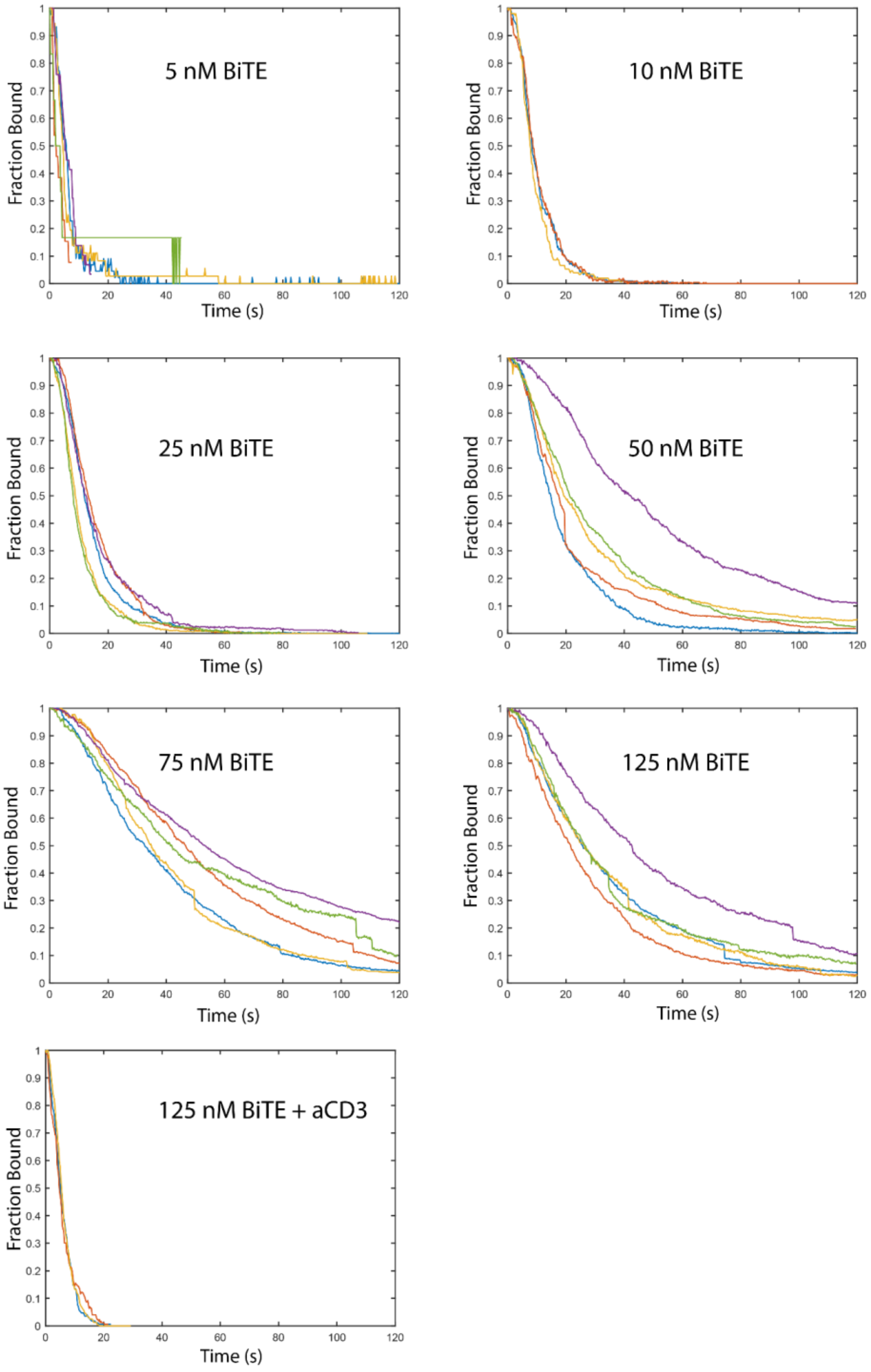

**Figure S4.** Individual trajectories for Jurkat cells on a BiTEs-coated surface with the indicated BiTE surface preparation. Total cells $N_{cells}$ = [128,576,2595,3005,4150,3492,584]

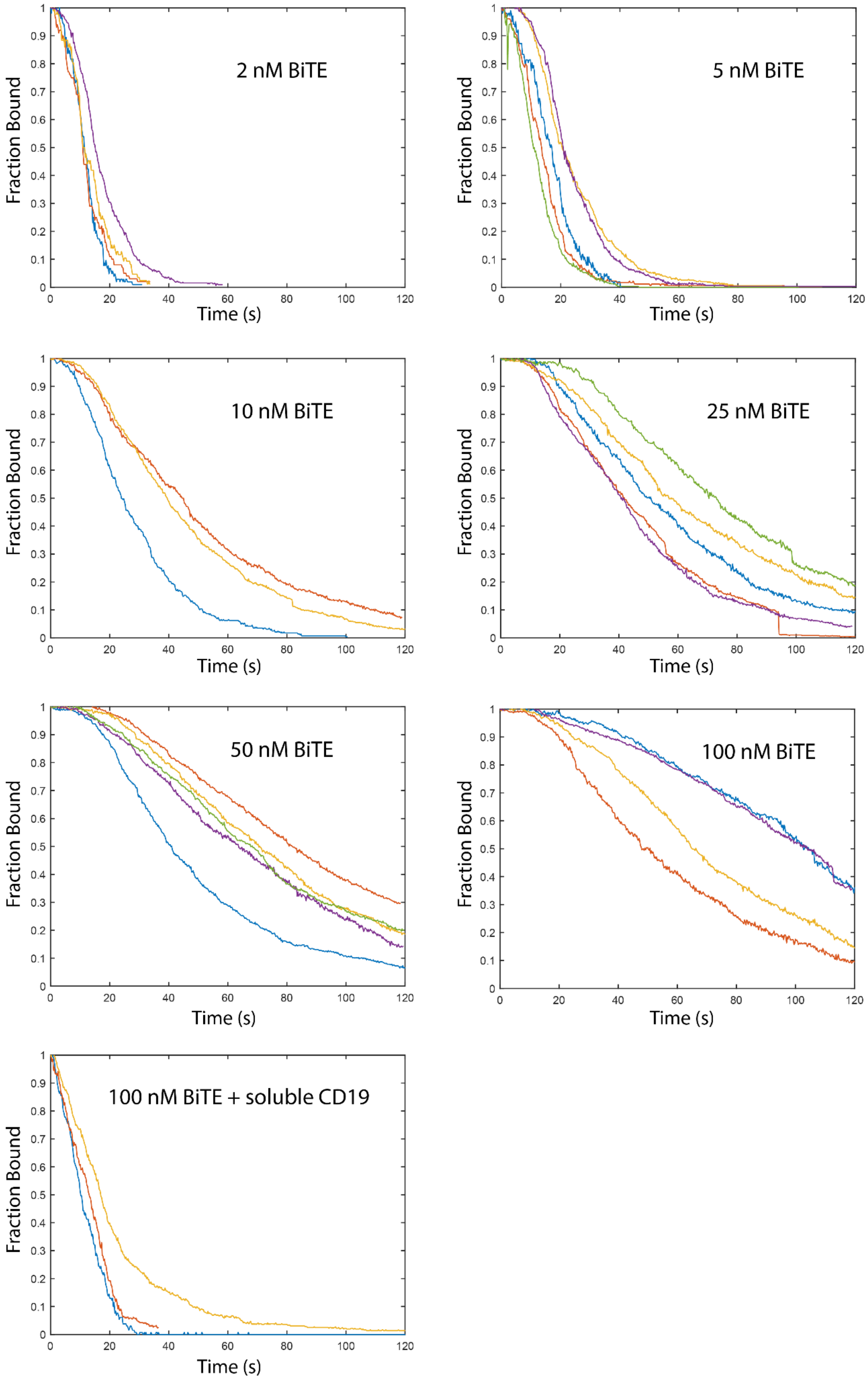

**Figure S5.** Individual trajectories for Nalm6 cells on a BiTEs-coated surface with the indicated BiTE surface preparation. Total cells $N_{cells}$ = [622, 1263, 1406, 2518, 3023, 2369,799].

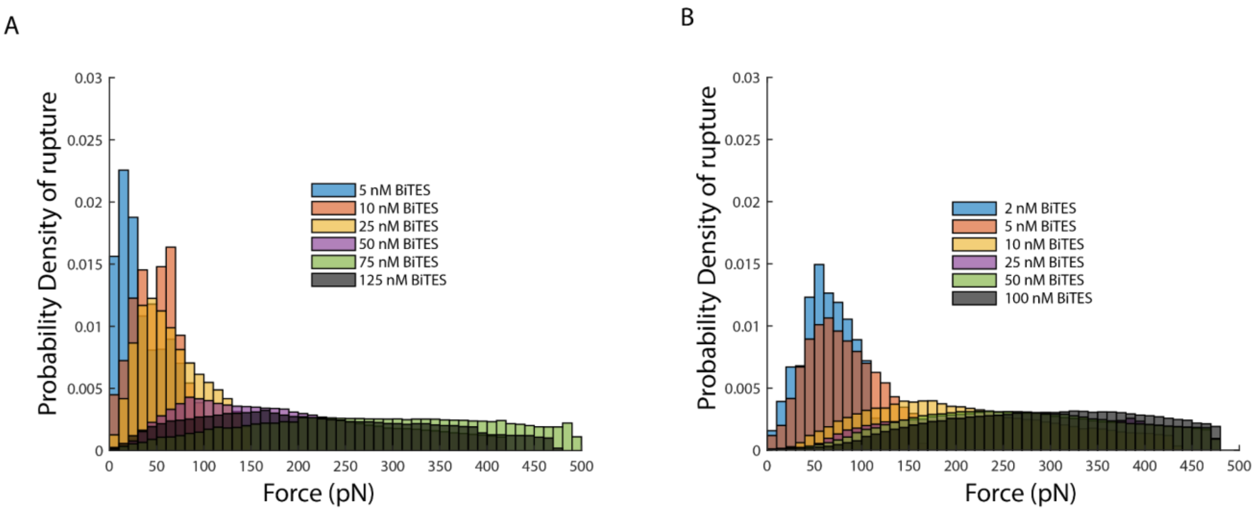

**Figure S6. A.** Rupture histogram for Jurkat cells binding to a BiTE functionalized surface prepared with a solution of the specified concentration. Total cells $N_{cells}$ = [128,576,2595,3005,4150,3492]. Fewer cells survive the 2-minute 1xg flip at lower concentrations, so fewer rupture events are observed. **B.** Rupture histogram for Nalm6 cells binding to a BiTE functionalized surface prepared with a solution of the specified concentration. Total cells $N_{cells}$ = [622, 1263, 1406, 2518, 3023, 2369]. The histograms are binned based on a nominal force calculation assuming constant cell density and size.

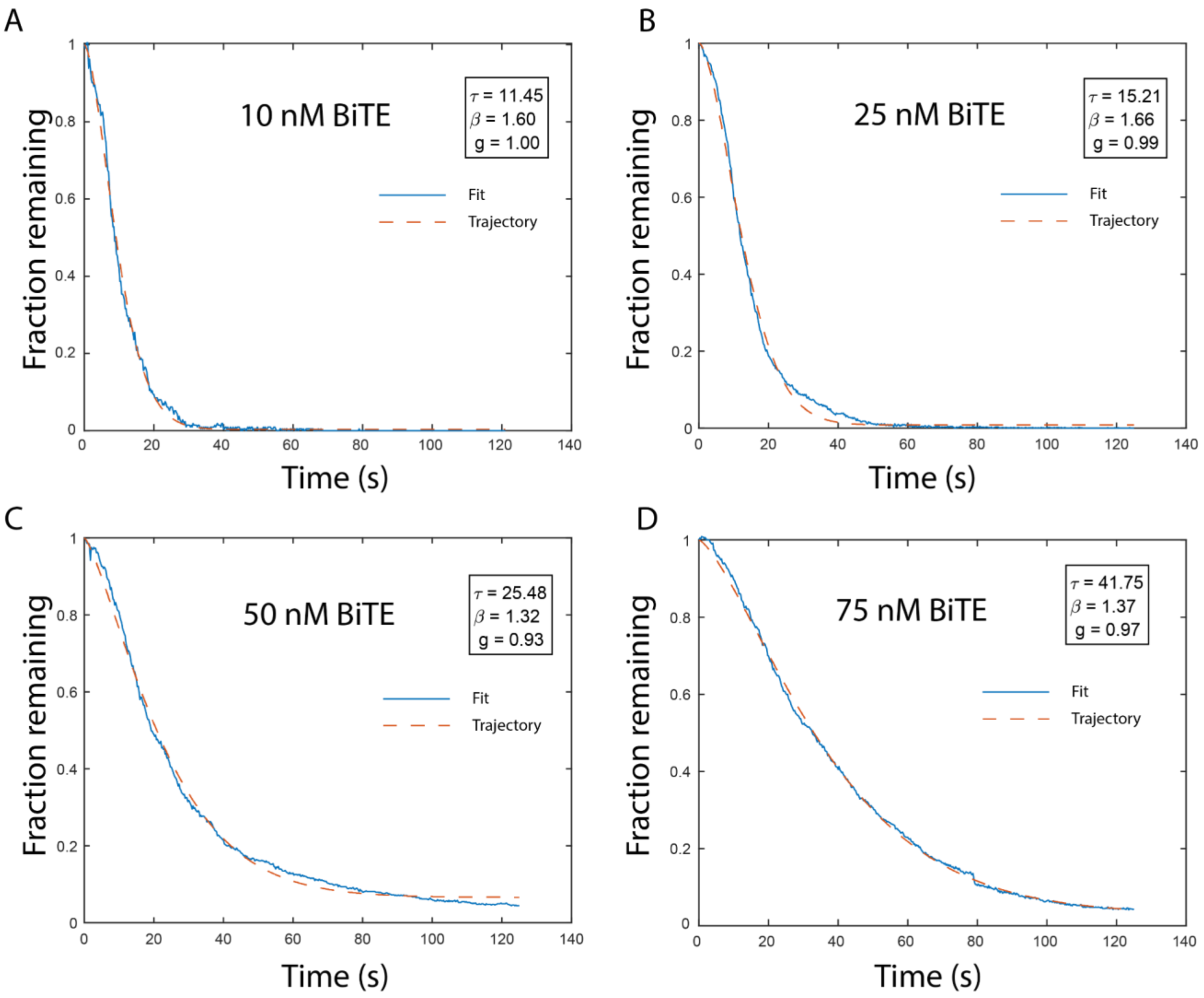

**Figure S7**. Example stretched exponential fits $1-g+g*exp(-(t/\tau)^{\beta})$. Trajectories are examples of Jurkat T-cell binding to BiTEs-functionalized surface prepared with the preparation concentration as indicated.

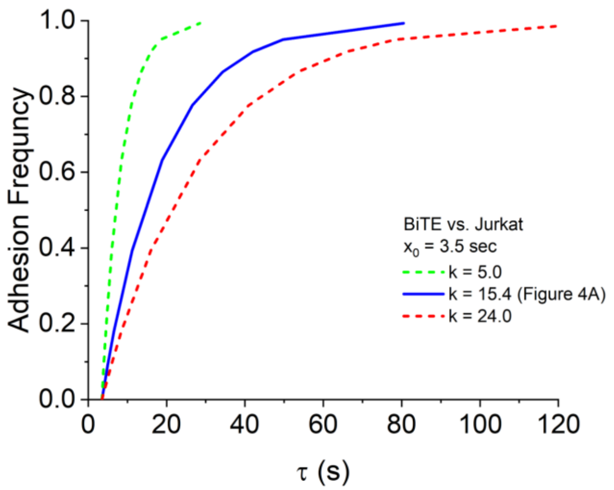

**Figure S8**. Examples of different parameter k values for the adhesion frequency and population lifetime relationship in Figure 4A (main text). The parametric equations (AF(λ) = 1 - CDF.Poisson (0,λ)) and population lifetime ($\boldsymbol{\tau}(\lambda) = k * \lambda + x_0$) were evaluated for different k values with $x_0$ set to 3.5. The increase of k value from 5.0 (green) to 24.0 (red) indicates shallower slope, i.e. a slower offrate under our force ramping conditions.

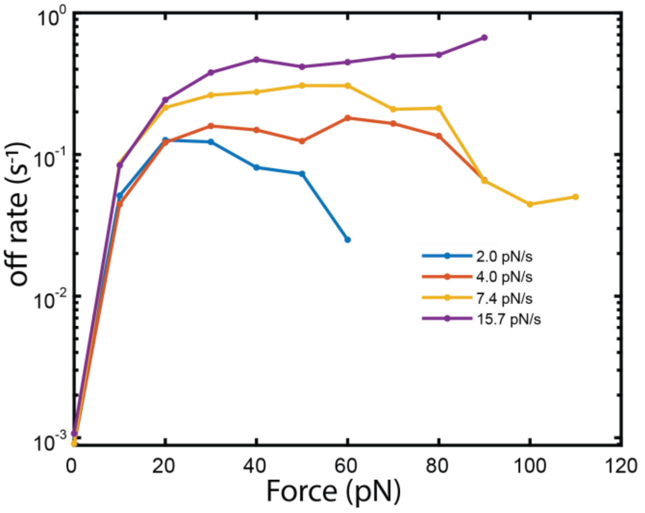

**Figure S9**. Jurkat cells binding to a surface prepared with 10 nM BiTE solution were measured with different loading rates (1-16 pN/s). The off rate at a given force was determined by normalizing the number of cells detaching at that force to the total number of available cells and scaling by the time interval [Evans 2009] [Dudko 2008]. The zero force off rate was calculated based off an interval during the flipping. Data listed in terms of increasing loading rate. $N_{trials}$ = [3,3,4,3], Total cells $N_{cells}$ = [633,606,1147,557]

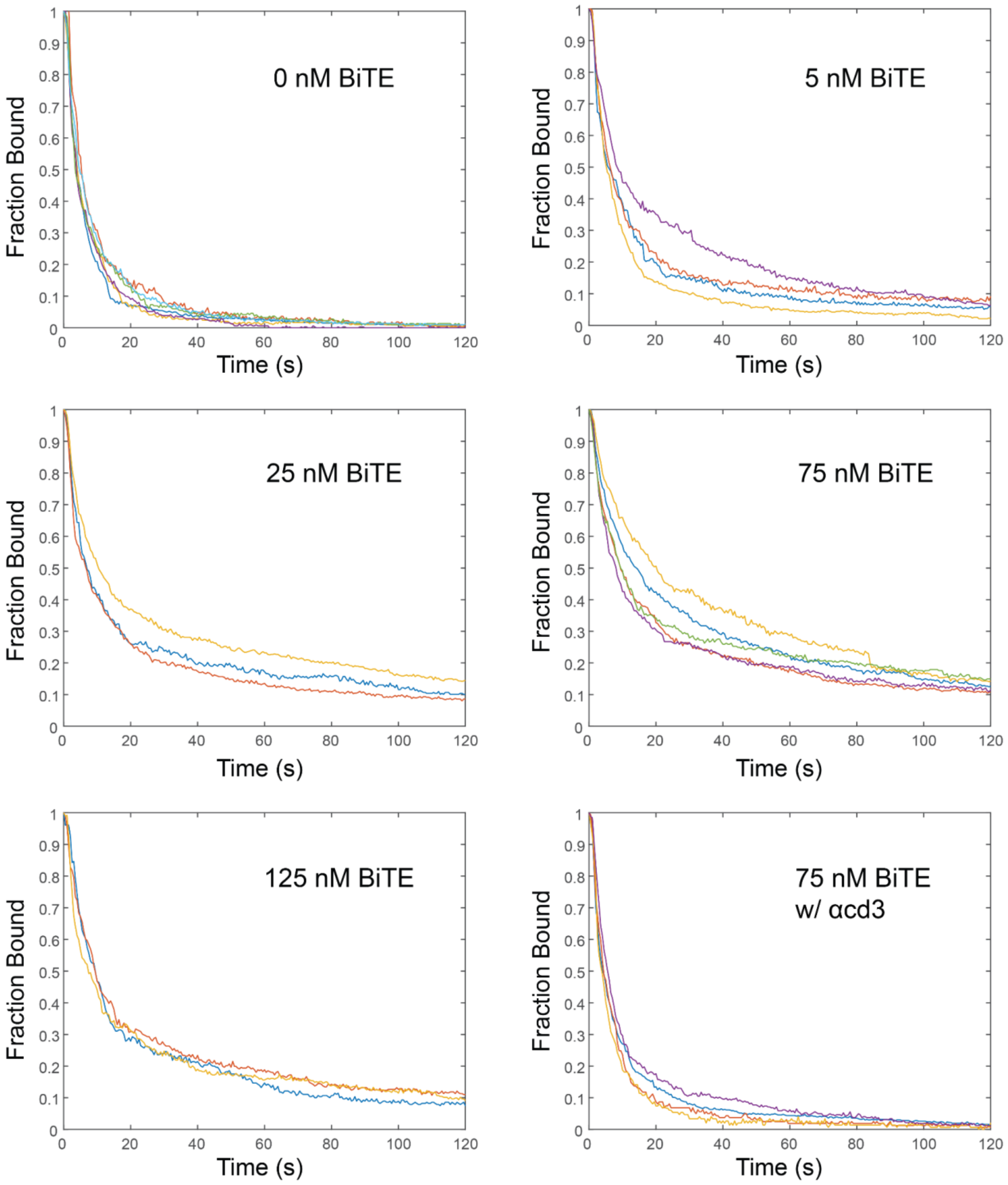

**Figure S10.** Individual trajectories of Jurkat cells binding to a Nalm6 monolayer at different BiTE concentrations with a 10 minute incubation. $N_{trials}$ = [6,4,3,4,5,3,4], Total cells $N_{cells}$ = [1946,1578, 1447, 1909, 1036, 1138]

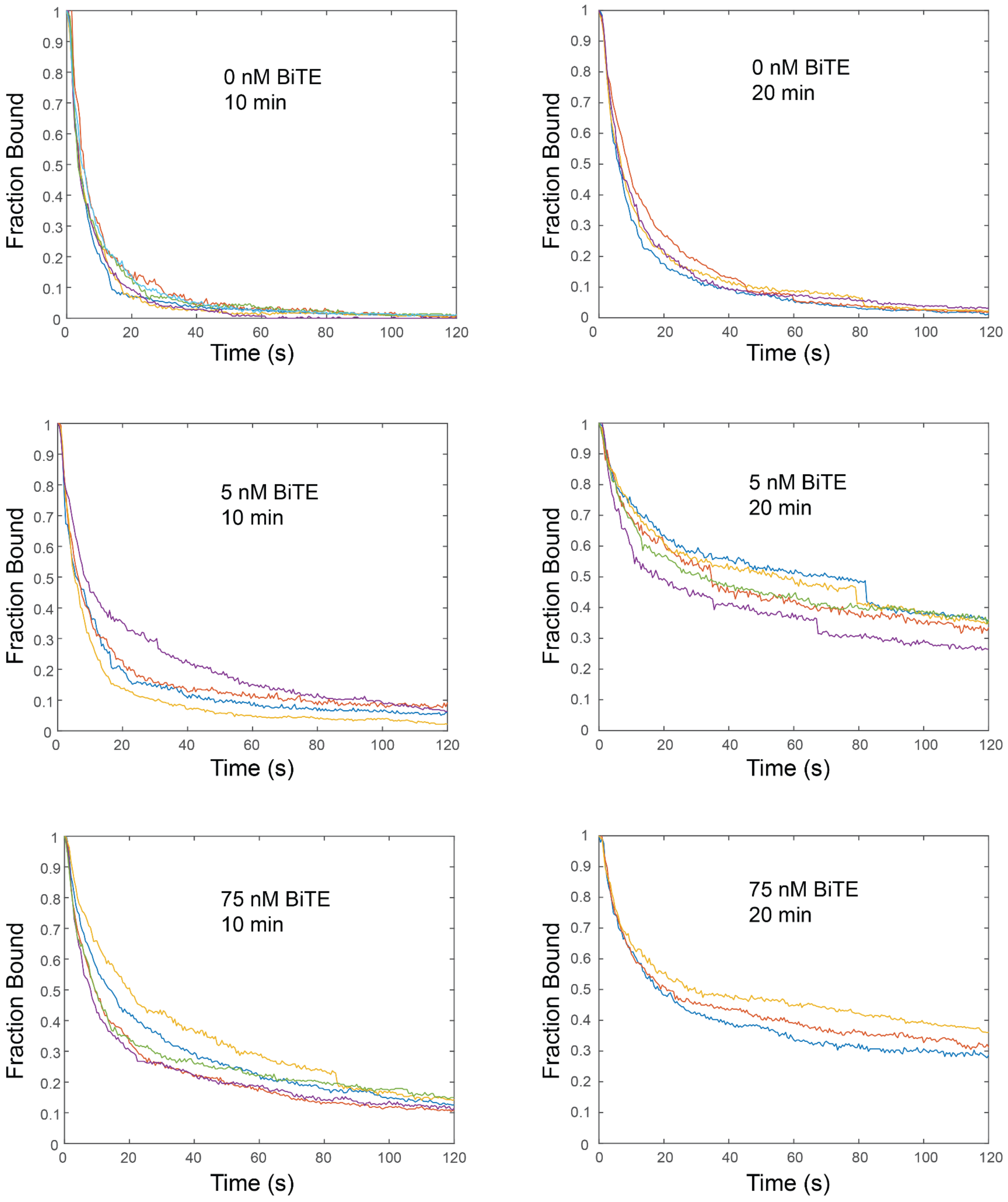

**Figure S11.** Individual trajectories of Jurkat cells binding to a Nalm6 monolayer with and without BiTE at different incubation times. Data listed: [0 nM 10 min, 5 nM 10 min, 75 nM 10 min, 0 nM 20 min, 5 nM 20 min,75 nM 20 min]. Number of trials: $N_{trials}$ = [6,4,5,4,5,3], Total cells $N_{cells}$ = [1946,1578, 1909, 1946, 2650,1319]

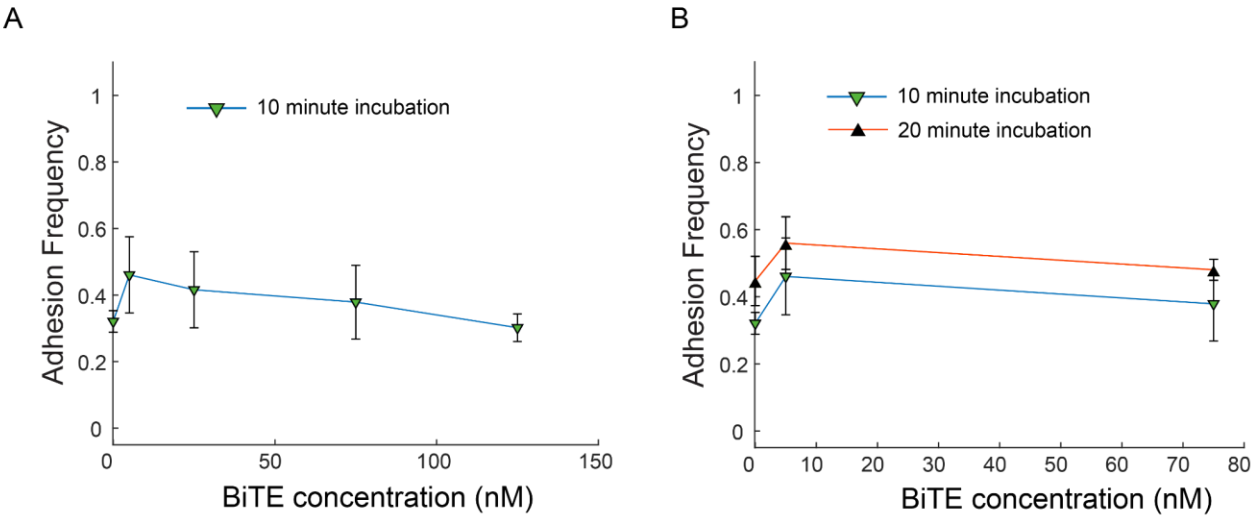

**Figure S12.** Fraction adhesion of Jurkat T-cell strength on BiTE treated Nalm6 cell monolayer. A) Adhesion frequency of Jurkat cells as a function of BiTE concentration used to treat the monolayer before attachment. No significant differences were observed over the concentration range tested. Error bars represent standard deviation over different trials. $N_{trials}$ = [6,4,3,5,3,4], $N_{cells}$ = [6011, 3473, 3485, 5238,3431, 5040] B) Adhesion frequency at two different attachment times and three BiTE concentrations. Error bars represent standard deviation over different trials. 20 minute incubation numbers: $N_{trials}$ = [4,5,3], Total cells $N_{cells}$ = [4427,4675,2703]

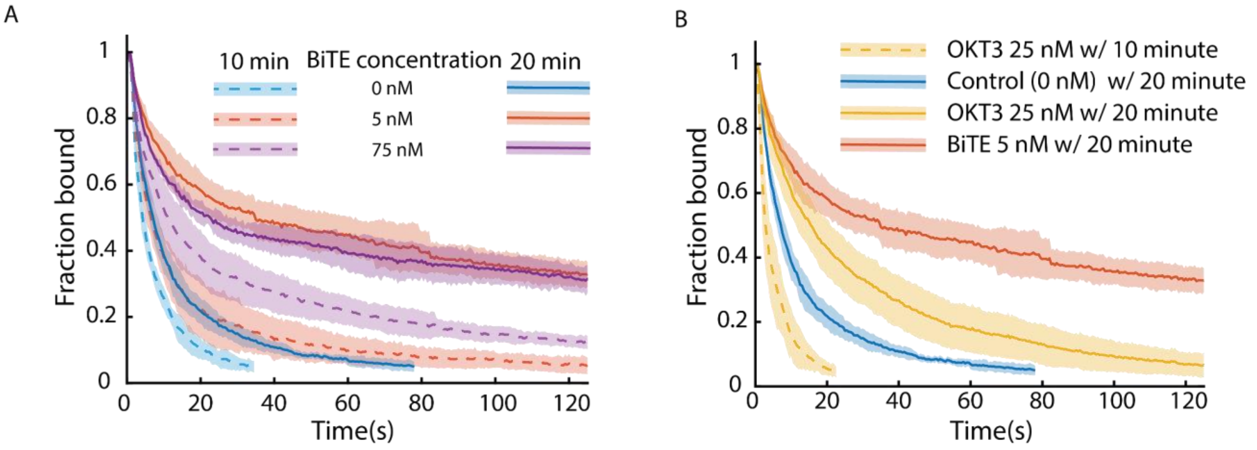

**Figure S13. A)** Time curves of Jurkat cells bound on Nalm6 monolayer at 10 and 20 minute attachment times and at three different BiTE concentrations (0 nM, 5nM, 75nM) with 4 pN/s force ramp. [0 nM 10 min, 5 nM 10 min, 75 nM 10 min, 0 nM 20 min, 5 nM 20 min,75 nM 20 min]. Number of trials: $N_{trials}$ = [6,4,5,4,5,3] $N_{cells}$ = [1946,1578, 1909, 1946, 2650,1319] **B)**Time curves of Jurkat cells bound on Nalm6 monolayer looking at the impact of incubation with anti-CD3 antibody OKT3 using different attachment times. OKT3 was supplied at 25 nM and compared to previous BiTES and control data. Data listed as [OKT3 25 nM 10 min, Control (0 nM), OKT3 25 nM 20 min, BiTE 5 nM 20 min]. Number of trials: $N_{trials}$ = [5,4,6,5] Total cells $N_{cells}$ = [704,1946,1914,2650]

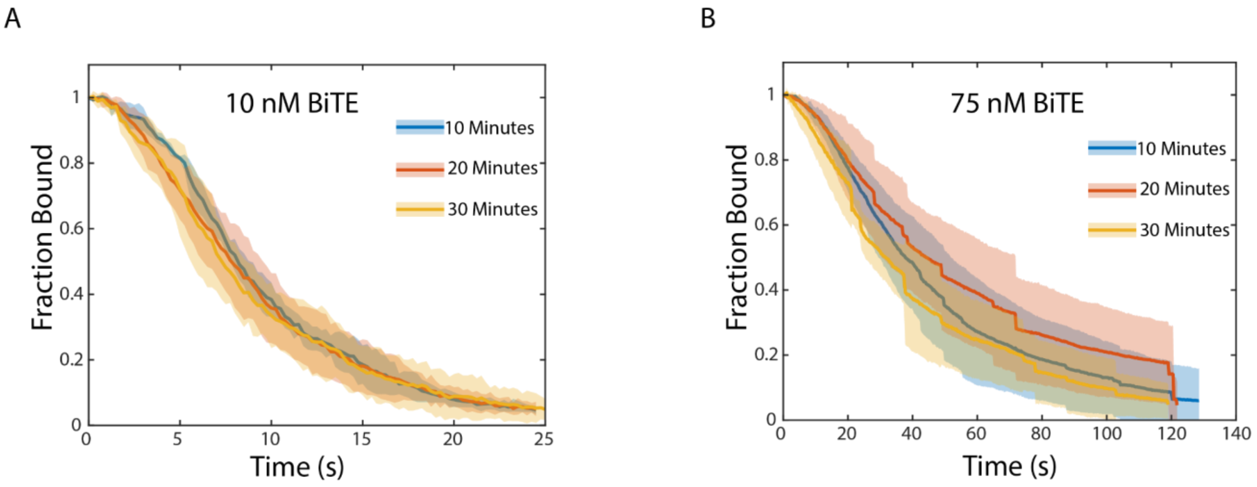

**Figure S14**. Ramping curves of Jurkat T-cells binding to a BiTE- functionalized surface with the specified BiTE surface concentration preparation measured using a 4 pN/s ramp. Cells were allowed to incubate on the surface for a different time, (10, 20, or 30 minutes). No difference was observed between the different incubation times for the cell-protein measurements. 10 nM BiTE - Number of trials: $N_{trials}$ = [3,3,3], $N_{cells}$ =[1791,2387,2544]. 75 nM BiTE - Number of trials: $N_{trials}$ =[4,5,3], Total cells $N_{cells}$ = [4163,4920,3546]

Table 1: Fluorescence Centrifuge Force Microscope Updated Parts list

| item # | Vendor | part # | Description | qty |
|---|---|---|---|---|
| 1 | Thorlabs | S05LEDM | SM05-Threaded Mount for LED | 1 |
| 2 | Thorlabs | SM1T1 | SM1 (1.035"-40) Coupler | 1 |
| 3 | Thorlabs | SM05RR | Retaining Ring for excitation filter | 1 |
| 4 | Thorlabs | SM05LTRR | Retaining Ring for excitation filter with rubber o-ring | 1 |
| 5 | Thorlabs | SM1LTRR | Retaining Ring for emission filter with rubber o-ring | 1 |
| 6 | Thorlabs | LEDRGBE | 627.5/525/467.5 nm Tri-Color LED | 1 |
| 7 | Thorlabs | SM1A6T | Diffuser and sample cell mount | 1 |
| 8 | SI Howard Glass Co | B-270 | Ø 25 mm, 0.9 mm Thick | 1 |
| 9 | Kapton Tape | PPTDE-3 | Sample Cell Assembly | 1 |
| 10 | Neuvitro | H-18-PLL | Coverslips with poly-L-lysine 18mm diameter #1 thickness | 1 |
| 11 | VWR | 63782-01 | Gold Seal, #1 19 mm coverglass | 1 |
| 12 | Thorlabs | SM1L03 | Sample Holder, SM1 Lens Tube, 0.3" Thread Depth | 1 |
| 13 | Thorlabs | SM1V05 | Focusing Ø1" SM1 Lens Tube | 1 |
| 14 | Edmund Optics | 86-815 | Olympus PLN 20X Objective, 0.40 NA, 1.2 mm WD | 1 |
| 15 | Thorlabs | SM1A3 | Objective Adaptor with External SM1 Threads and Internal RMS Threads | 1 |
| 16 | Thorlabs | AC254-100 | Tube Len, f=100.0 mm, Ø1" Achromatic Doublet, ARC: 400-700 nm | 1 |
| 17 | Thorlabs | SM1RR | Tube Lense SM1 Retaining Ring | 2 |
| 18 | Thorlabs | SM1M20 | Objective SM1 Lens Tube Without External Threads, 2" Long | 1 |
| 19 | Thorlabs | SM1A6T | Adaptor with External SM1 Threads and Internal SM05 Threads, 0.40" Thick | 2 |
| 20 | | | Custom made turning block, aluminum | 1 |
| 21 | Thorlabs | PFE10-P01 | Turning Mirror, 1" Silver Elliptical Mirror, 450 nm - 20 µm | 2 |
| 22 | Thorlabs | SM1NT | Camera SM1 (1.035"-40) Locking Ring, Ø1.25" Outer Diameter | 1 |
| 23 | Thorlabs | SM1A9 | Camera Adaptor with External Cmount Threads and Internal SM1 Threads | 1 |
| 24 | Teledyne FLIR | BFS-PGE-88S6M-C | Sony IMX267 CMOS sensor, 4096 x 2160 resolution, 3.45x3.45 um pixel size | 1 |
| 25 | IMC Network | 855-10734 | MiniMc-Gigabit Twisted Pair to Fiber Media Converter | 1 |
| 26 | PrinceTel | MJX | Fiber Optic Rotary Joint | 1 |

| | | | | |
|---|---|---|---|---|
| 27 | Chroma | 89402x | Multiband pass excitation filter, unmounted, 12.5 mm diameter | 1 |
| 28 | Chroma | 89402m | Multiband pass emission filter, unmounted, 25 mm diameter | 1 |
| 29 | TT Electronics | OPB732 | Infrard LED and Phototransistor Long Distance Reflective Switch for RPM measurement | |
| 30 | National Instruments | NI USB-6008 | DAQ board used to connect photo switch and computer for rpm measurement | 1 |
| 31 | Adafruit | 1903 | PowerBoost 500 Basic - 5V USB Boost @ 500 mA from 1.8V+ | 2 |
| 32 | Adafruit | 3500 | Adafruit Trinket M0 microcontroller | 1 |
| 33 | Amazon | | Batteries 2500 mAh TR 14500 3.7v Li-ion | 6 |
| | | Other | 3D printed battery pack, centrifuge bucket, wires | |

**Table 2:**

| **Spec.** | **CFM** |
|---|---|
| Applied force/cell | 0 ~ 500 pN* (3,000 rpm) |
| Linear loading rate | Up to ~20 pN/s |
| Measurement cells #<br>(in the field of view) | ~1,000 cells/run |
| Fluorescent image | Up to 2 fluorescent colors |
| Image resolution | 170 x 170 nm/pixel |
| Frame speed | 8 frames/sec |
| Sample chamber | Disposable |

*Depending on the cell size and density
